# Proteomic and functional analyses of the periodic membrane skeleton in neurons

**DOI:** 10.1101/2020.12.23.424206

**Authors:** Ruobo Zhou, Boran Han, Roberta Nowak, Yunzhe Lu, Evan Heller, Chenglong Xia, Athar H. Chishti, Velia M. Fowler, Xiaowei Zhuang

## Abstract

Actin, spectrin, and associated molecules form a membrane-associated periodic skeleton (MPS) in neurons. The molecular composition and functions of the MPS remain incompletely understood. Here, using co-immunoprecipitation and mass spectrometry, we identified hundreds of candidate MPS-interacting proteins that span diverse functional categories. We validated representative proteins in several of these categories, including previously unknown MPS structural components, as well as motor proteins, cell adhesion molecules, ion channels, and signaling proteins, demonstrating periodic distributions of ∼20 proteins in neurons using super-resolution imaging. Genetic perturbations of the MPS and its interacting proteins further suggested functional roles of the MPS in axon-axon and axon-dendrite interactions and in axon diameter regulation, and implicated the involvement of MPS interactions with cell adhesion molecules and non-muscle myosin in these roles. These results provide new insights into the interactome of the MPS, and suggest new functions of the MPS in neurons.

## Introduction

Super-resolution imaging has recently revealed a membrane-associated periodic skeleton (MPS) in neurons^1^. The neuronal MPS contains some structural components that are homologous to the membrane skeleton of erythrocytes^2, 3^, but adopts a distinct ultrastructure in neurons, in which actin filaments are assembled into ring-like structures, and the adjacent actin rings are connected by spectrin tetramers consisting of two αII-spectrin and two β-spectrin (βII, βIII or βIV) subunits, forming a one-dimensional (1D) periodic structure with ∼190-nm periodicity underneath the plasma membrane of neurites^1^. In mature neurons, the MPS structure spans the entire axonal shaft, including both the axon initial segment and distal axon^1, 4, 5^, and has also been observed in a substantial fraction of dendritic regions^4, 6, 7^. In soma and a fraction of dendritic regions, a 2D polygonal MPS structure resembling the membrane skeleton of erythrocytes has been observed^7^, but the average end-to-end distance of the spectrin tetramer connecting the two adjacent actin nodes in the native environment is significantly greater in neurons than in erythrocytes^7, 8^. A highly prevalent structure in the nervous system, the MPS is present in diverse neuronal types, including excitatory and inhibitory neurons in both central and peripheral nervous systems^9, 10^, and across diverse animal species, ranging from C elegans to humans^10^. This submembrane lattice structure can organize transmembrane proteins^1, 4, 6, 11-13^, and it has been shown recently that the MPS can function as a signaling platform that coordinates the interactions of key signaling proteins and enables signal transduction in neurons^13^. Additional functional roles of the MPS have also been implicated in axon stability under mechanical stress^1, 14^, mechanosensation^15^, diffusion restriction at the AIS^16^, and axon degeneration^17, 18^. Disruption of the MPS causes a range of neurological impairments in mice^19, 20^.

Despite the importance of the MPS in neurons, its molecular components and interacting partners have not been systemically investigated, which limits our understanding of the functions of the MPS and the molecular mechanisms underlying these functions. In addition to actin and spectrin, two other structural components of the MPS have been identified: adducin, a protein that caps the fast-growing end of actin filaments, and ankyrin, an adaptor protein that can anchor transmembrane proteins to the MPS^1, 4, 5^. A few transmembrane proteins have also been observed to associate with the MPS structures in the AIS region, including a couple of ion channels and adhesion molecules^1, 4, 6, 11, 12^. Recently, it has been shown that several transmembrane signaling proteins, including the G-protein coupled receptor CB1, cell adhesion molecule NCAM1, and two receptor tyrosine kinases (RTKs), TrkB and FGFR, can be recruited to the MPS structure in response to extracellular stimuli to enable RTK transactivation in neurons^13^. The list of MPS components and interacting partners is, however, likely far from complete.

In this work, we used co-immunoprecipitation (co-IP) to pull down MPS-associated proteins from the adult mouse brain tissue and cultured neurons, followed by mass spectrometry for a proteomic-scale identification of the candidate structural components and interacting partners of the MPS. This analysis revealed hundreds of potential candidate MPS-interacting proteins, which span many functional categories and cellular pathways. In addition, we used quantitative mass spectrometry to investigate how the expression levels of proteins are differentially regulated at the proteomic scale upon disruption of the MPS. We further used stochastic optical reconstruction microscopy (STORM)^21, 22^, a super-resolution imaging method, to validate the MPS association for a subset of the candidate proteins in several important functional categories, including actin-binding proteins, motor proteins, cell adhesion molecules, ion channels, and other signal transduction proteins. Using genetic knockout or shRNA knockdown, we identified several structural molecules that are essential for the MPS formation. Our genetic perturbation experiments further suggested functional roles of the MPS in axon-axon and axon-dendrite interactions, as well as in axon diameter regulation.

## Results

### Proteomic analysis of candidate structural components and interacting partners of the MPS

We used co-IP and mass spectrometry to pull down and identify potential candidate structural components and interacting partners of the MPS in neurons (Fig. 1a). Magnetic beads were coated with the antibody that can specifically bind to a bait protein previously known to be an MPS structural component, such as βII-spectrin, αII-spectrin and α-adducin^1^, and these beads were then incubated with the lysate from either adult mouse brains or cultured mouse hippocampal neurons at 20 days in vitro (DIV) to allow capturing of the proteins that bind directly or indirectly to these known MPS components. The co-immunoprecipitated protein mixtures were subsequently analyzed using SDS-PAGE and mass spectrometry.

**Fig. 1:**
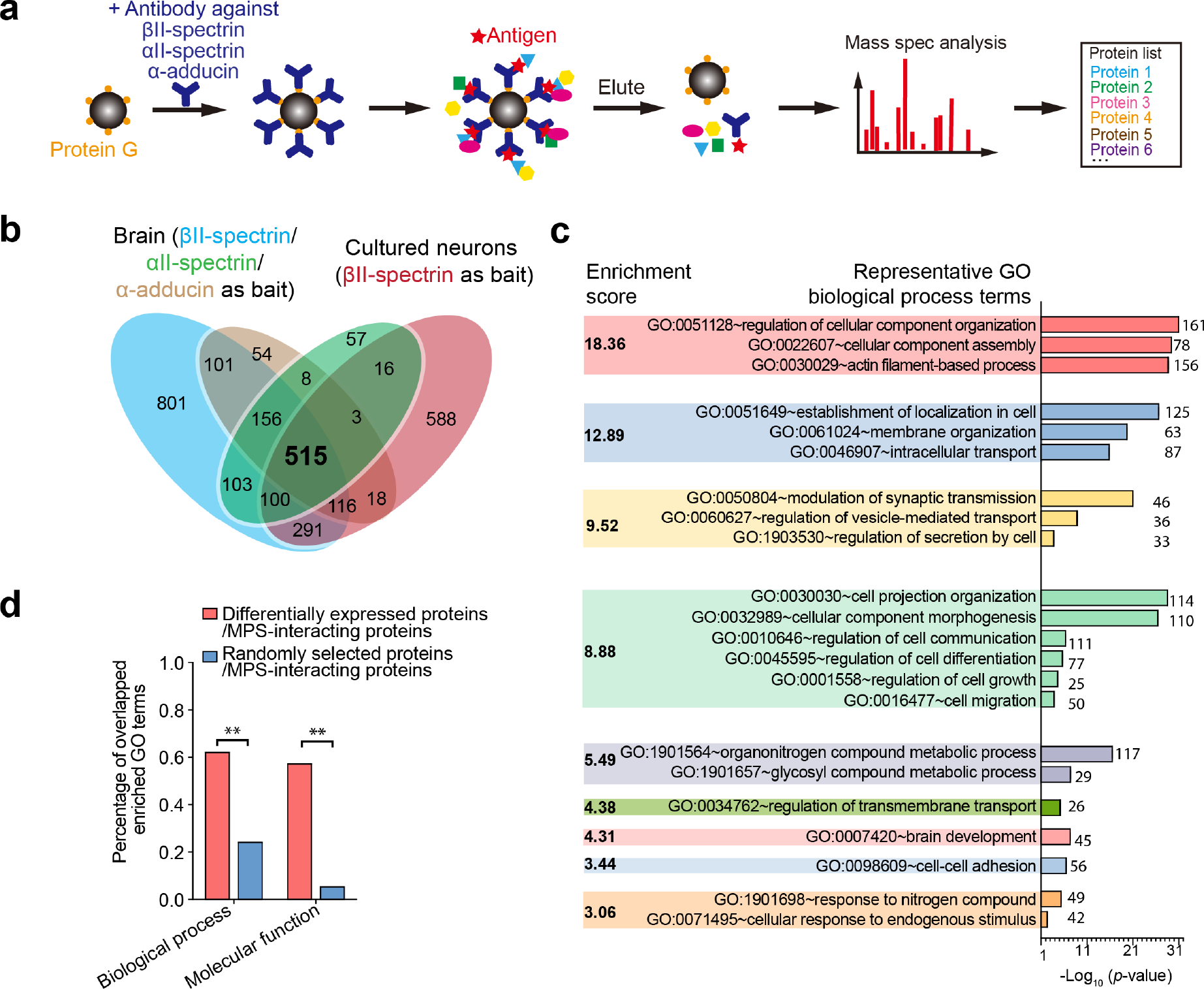
Proteomic analysis and GO term enrichment analysis of candidate MPS-interacting proteins. **a** Schematic for co-immunoprecipitation (co-IP) based mass spectrometry identification of candidate MPS-interacting proteins. The antibody against a bait protein, such as βII-spectrin, αII-spectrin, and α-adducin, was attached to protein-G-coated beads. The antibody-coated beads were then used to capture the bait protein and the co-immunoprecipitated proteins from the adult mouse brain lysate or the cultured hippocampal neuron lysate. The co-immunoprecipitated proteins were identified using mass spectrometry. **b** Venn diagram showing the overlap of the identified proteins in four co-IP experiments, including the co-IP experiments from the mouse brain using βII-spectrin (cyan), αII-spectrin (green) or α-adducin (brown) as the bait, and the co-IP experiment from cultured hippocampal neurons (DIV 20) using βII-spectrin as the bait (red). Proteins that were commonly identified in all of the four co-IP experiments were considered candidate MPS-interacting proteins in the following analyses. **c** Functional annotation clustering of the enriched Gene Ontology (GO) terms in the biological process (BP) category for the 515 candidate MPS-interacting proteins. Enriched GO BP terms were generated and clustered using the DAVID 6.8 platform with the enrichment score listed for each color-coded GO term cluster. Representative GO BP terms for each GO term cluster were shown. The corresponding *p*-values (bars) and protein numbers (next to the bars) for each enriched GO BP term are shown on the right. **d** Red: Percentages of the enriched GO terms of the candidate MPS-interacting proteins that overlap with the enriched GO terms of the 1204 differentially expressed proteins in βII-spectrin knockdown neurons. Blue: Percentages of the enriched GO terms of the candidate MPS-interacting proteins that overlap with the enriched GO terms of 1204 randomly selected genes from the mouse genome. The 1204 differentially expressed proteins, with fold change >1.2 (for up regulation) or < 0.8 (for down regulation) in the βII-spectrin knockdown neurons as compared to the neurons treated with scrambled (control) shRNA, were determined using the DESeq2 package^68^ to test the statistical significance for up and down regulation of each protein (*p* < 0.01). ** indicates *p* < 0.005 (Chi-squared test).

The co-immunoprecipitated proteins from either adult mouse brain or cultured neurons showed notable enrichment of spectrin and actin, as well as other proteins, at the expected molecular weights in the SDS-PAGE gel images, whereas co-IP using beads coated with a control IgG hardly pulled down anything (Supplementary Fig. 1a, b). Furthermore, treatment with actin disrupting drugs (LatA and CytoD), which are known to disrupt the MPS^1, 4, 7^, substantially reduced the amount of co-immunoprecipitated proteins (Supplementary Fig. 1a, b).

Next, we used mass spectrometry to identify the co-immunoprecipitated proteins. To reduce the number of potential false positives of the identified MPS structural components and interacting proteins, we considered only the 515 proteins commonly identified in all four co-IP experiments –– i) using βII-spectrin, as the bait in brain tissues, ii) using αII-spectrin as the bait in brain tissues, i) using α-adducin as the bait in brain tissues, and iv) using βII-spectrin as the bait in cultured neurons (Fig. 1b), although this criterion may filter out some of the true MPS-interacting proteins. Actin, βII-spectrin, α-adducin and ankyrin, which have been previously shown to be structural components of the neuronal MPS^1^, are indeed among the 515 proteins identified here. It is worth noting that because the lysates used in the co-IP experiments were not specific to the fraction of any specific neuronal sub-compartment, the identified candidate protein list could thus include structural components and interacting partners of the MPS structures, for both 1D and 2D forms, in axons, dendrites, and soma. As a cautionary note, as is typically true for the proteomes identified by co-IP and mass spectrometry, the proteins included in this list are potential candidates of structural components and interacting partners of the MPS that need to be further validated, as we did for a subset of these candidates shown in later sections.

For global analysis of these identified candidate proteins, we performed gene ontology (GO) term analysis of the biological process (BP) category using the DAVID platform^23^. The functional annotation clusters of enriched GO BP terms (*p* value < 0.05) showed potential relationship of the MPS to many biological processes (Fig. 1c), including actin filament-based process, membrane organization, vesicle-mediated transport, synaptic transmission, cell morphogenesis, cell differentiation and growth, cell communication, transmembrane transport, cell adhesion and migration, brain development, etc. Similar GO term analysis of the molecular function (MF) category also suggested potentially involvement of the MPS in diverse cellular functions (Supplementary Fig. 1c).

Next, we used multiplexed quantitative mass spectrometry based on Tandem Mass Tag (TMT) isobaric labeling^24^ to systematically determine the protein abundance changes in the cultured neurons upon βII-spectrin knockdown, which is known to disrupt the MPS^4, 7^. Among the proteins detected in our quantitative mass spectrometry experiment, 1204 showed statistically significant up or down regulation in βII-spectrin knockdown neurons as compared to control neurons treated with scrambled shRNA. 468 of the 515 candidate MPS-interacting proteins identified above were detected by our quantitative mass spectrometry experiment, and among these 162 were found significantly up- or down-regulated in the βII-spectrin knockdown neurons. Notably, although we do not expect all MPS-interacting proteins to be up or downregulated upon MPS disruption, nor do we expect all differentially expressed proteins to necessarily interact with the MPS, we found that the enriched GO terms of the 515 candidate MPS-interacting proteins determined by co-IP and mass spectrometry largely overlapped with the enriched GO terms of the 1204 differentially expressed proteins upon MPS disruption (Fig. 1d and Supplementary Fig. 2), suggesting that proteins probed in these two experiments likely function in common or related biological processes.

### Validation of candidate MPS components using super-resolution imaging

The co-IP-based mass spectrometry analysis provided a list of candidate MPS-interacting proteins, not all of which are necessarily associated with the MPS. Additional experiments are needed to validate the association of these proteins with the MPS. Among the identified candidate MPS-interacting proteins, a notable number of them were actin-binding proteins, which potentially bind to the actin filaments in the MPS and hence regulate the stability of the MPS. To validate these actin-binding MPS components, we imaged some of them using STORM^21, 22^ to examine whether these proteins exhibit periodic distributions along axons, in a way similar to actin, spectrin and α-adducin as previously reported^1^.

To visualize these proteins, we used either immunostaining with antibody against the endogenous proteins, or moderate expression of GFP-fusion proteins through low-titer lentiviral transfection followed by immunolabeling using anti-GFP antibody. For the molecules that have high-quality antibodies, the first approach generally provided higher quality images, as indicated by a higher degree of periodicity of the protein distribution along axons for MPS structural components (Supplementary Fig. 3a), potentially because the GFP-fusion proteins cannot be as efficiently incorporated into the MPS due to perturbation of the GFP tag, overexpression, or competition with endogenous proteins. Indeed, expressing GFP-fusion proteins in neurons where the endogenous proteins were depleted can improve the incorporation efficiency of the fusion proteins into the MPS (Supplementary Fig. 3b). It is also worth noting that antibody quality for immunolabeling can vary substantially, even for the same target protein (Supplementary Fig. 3c). We therefore typically screened multiple antibodies to identify the highest quality antibody for imaging and when antibodies with adequate quality did not exist, we used the GFP-fusion approach.

Using these labeling strategies, we imaged five actin-binding proteins, including αII-spectrin, tropomodulin 1, tropomodulin 2, dematin, and coronin 2B. In the STORM images, αII-spectrin, tropomodulin 1, tropomodulin 2, dematin, and coronin 2B exhibited periodic distributions along the axons (Fig. 2a-e). To quantify the periodic distribution, we calculated the average 1D autocorrelation function and its amplitude from randomly selected axon regions for each imaged protein molecule. We determined the amplitude of the autocorrelation function as the difference between the value at the first 190-nm peak and the average value at the first two valleys at ∼95 nm and ∼285 nm, which provides a quantitative measure of the periodicity of the measured distribution. We found that all five examined actin-binding proteins showed periodic autocorrelation with a periodicity of ∼190-nm but with varying amplitudes (Fig. 2a-e). These results suggest several new structural components for the MPS, in addition to the previously identified components, actin, spectrin, α-adducin, and ankyrin. Among these proteins, tropomodulin 1 and dematin are also structural components of the erythrocyte membrane skeleton, whereas tropomodulin 2 and coronin 2B are not found in the erythrocyte membrane skeleton^2, 3^.

**Fig. 2:**
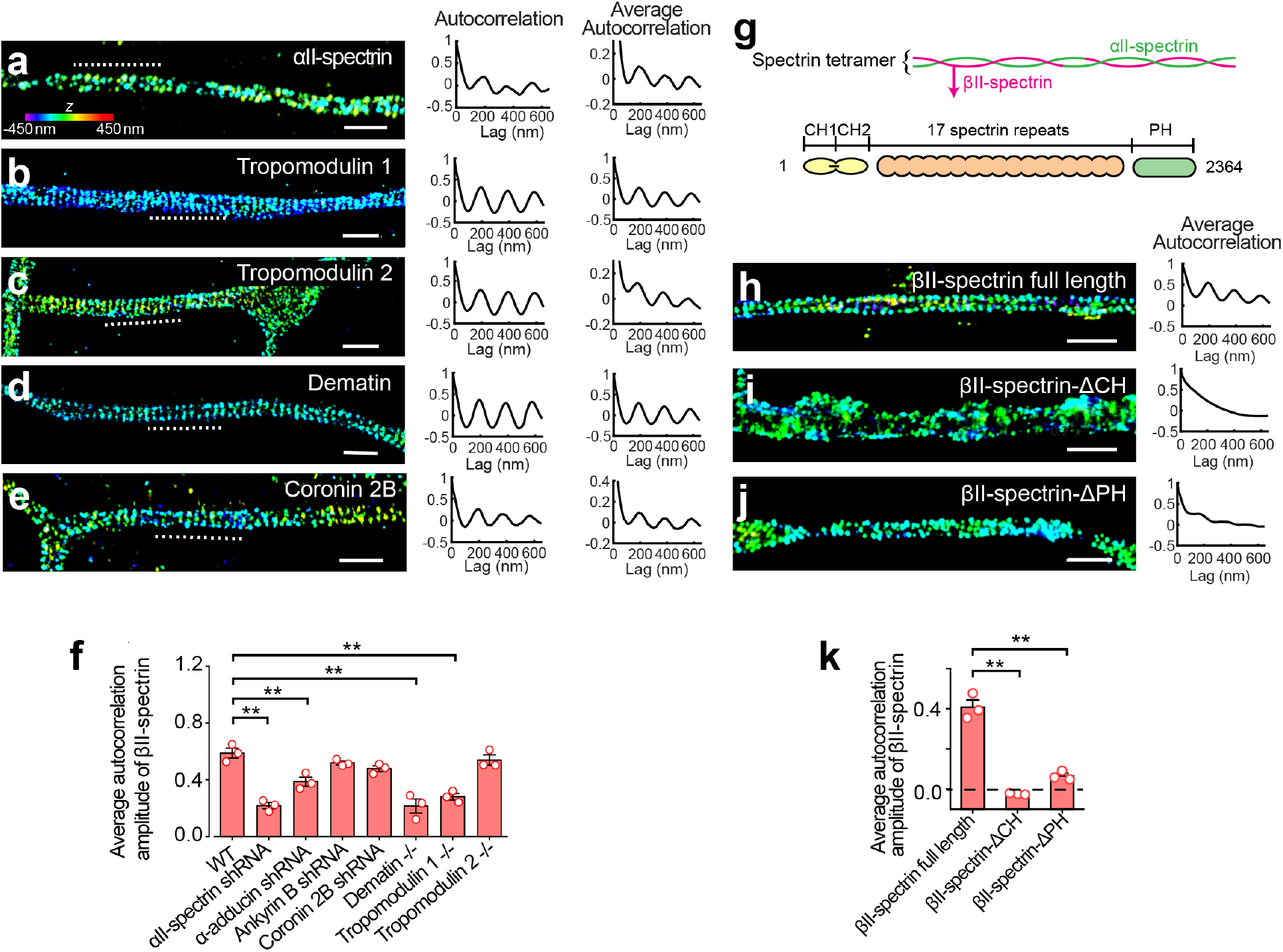
Validation of actin-binding MPS components and identification of essential structural component of the MPS using super-resolution imaging. **a-e** Left: 3D STORM images of axon regions of cultured hippocampal neurons immunostained for αII-spectrin (**a**), tropomodulin 1 (**b**), tropomodulin 2 (**c**), dematin (**d**), and coronin 2B (**e**). Middle: One-dimensional (1D) autocorrelation functions of the imaged molecules for the axon region indicated by the dashed line in the left panels. Signals were projected to the longitudinal axis of the axon segment, and the 1D autocorrelation functions were calculated using the projected signals. Right: Average 1D autocorrelation functions of the imaged molecules over many randomly selected axon regions. αII-spectrin, dematin and coronin 2B were immunolabeled with their corresponding antibodies. Tropomodulin 1 and tropomodulin 2 were labeled using moderate expression of GFP-tagged proteins through low-titer lentiviral transfection, which were in turn immunolabeled using anti-GFP antibody. GFP-tagged tropomodulin 1 was expressed in tropomodulin 1 knockout neurons; GFP-tagged tropomodulin 2 was expressed in WT neurons. Scale bars: 1 μm. Colored scale bar indicates the z-coordinate information. **f** Average 1D auto-correlation amplitudes of the βII-spectrin distribution, indicating the degree of periodicity of the MPS, calculated from many axon segments of untreated neurons, neurons transfected with adenoviruses expressing αII-spectrin shRNA, α-adducin shRNA, ankyrin B shRNA, or coronin 2B shRNA, and neurons cultured from tropomodulin 1, tropomodulin 2 or dematin knockout mice. The average 1D auto-correlation amplitude was defined as the difference between the first peak (at ∼190 nm) and the average of the two first valleys (at ∼95 nm and ∼285 nm, respectively) of the average 1D auto-correlation function^4^. **g** Domain organization of βII-spectrin. **h-j** Left: 3D STORM image of axon regions of βII-spectrin knockout neurons transfected with plasmid expressing GFP-tagged full length βII-spectrin (**h**), βII-spectrin-ΔCH mutant (**i**) or βII-spectrin-ΔPH mutant (**j**). The proteins were visualized by immunostained with anti-GFP antibody. Scale bars: 1 μm. Right: average 1D autocorrelation functions of the imaged molecules indicated in the left panels. **k** Average 1D auto-correlation amplitudes for the distribution of GFP-tagged full length βII-spectrin and the distributions of the two GFP-tagged βII-spectrin truncation mutants, as described in (**h-j**), calculated from many GFP-positive axon segments. Data are mean ± s.e.m. (*n* = 3 biological replicates for each condition; for each biological replicate, 50-100 axonal regions were examined). * indicates *p* < 0.05 and ** indicates *p* < 0.005 (unpaired Student’s t-test).

### Proteins essential for the MPS formation or maintenance

To test whether these structural molecules are essential for MPS formation and/or maintenance in neurons, we depleted these proteins using either shRNA knockdown (Supplementary Fig. 4a) or knockout mice, and used STORM imaging to quantify the integrity of the MPS under each genetic perturbation condition by measuring the degree of periodicity of the βII-spectrin distribution in axons, as indicated by the average autocorrelation function amplitude. Compared to the wild type (WT) neurons, the MPS was disrupted or partially disrupted in the neurons depleted of αII-spectrin, α-adducin, tropomodulin 1, or dematin, whereas depletion of ankyrin B, coronin 2B, or tropomodulin 2 did not significantly disrupt the MPS (Fig. 2f and Supplementary Fig. 4b). Our observation that αII-spectrin is essential for the MPS formation in axons is consistent with our previous results that the MPS is disrupted in βII-spectrin knockdown neurons^4, 7^ and the previous result that αII-spectrin knockout disrupts the MPS in the AIS region^19^. Our observation that the MPS is partially disrupted in the α-adducin depleted mouse neurons is consistent with the previous result reported in *Drosophila* neurons carrying the *hts*^*1*^ null mutant allele, which abolishes *Drosophila* adducin^25^. It has been also reported that the axons in α-adducin knockout mouse neurons showed enlarged diameters while the MPS was not fully disrupted, although the level of disruption was not quantitatively assessed in this previous study^26^.

In addition to examining which structural proteins are essential for the MPS formation/maintenance, for βII-spectrin, a known key component of the MPS^4, 7^, we further examined the role of its various protein domains. βII-spectrin contains the N-terminal CH domain responsible for actin binding and the C-terminal pleckstrin homology (PH) domain responsible for binding to PI(4,5)P_2_ phosphatidylinositol lipids as well as some membrane proteins (Fig. 2g)^2^. We thus examined the effect of deleting the CH or PH domain on the MPS integrity. We expressed GFP-tagged full length βII-spectrin, or GFP-tagged βII-spectrin mutants with either CH or PH domain deleted (βII-spectrin-ΔCH or βII-spectrin-ΔPH), in neurons cultured from brain-specific βII-spectrin knockout mice, which we generated by crossing *βII-Spec*^*flox/flox*^ mice^27^ with the Nestin-Cre mouse line. We found that only the full length βII-spectrin could support the MPS formation, exhibiting a periodic distribution along axons, whereas βII-spectrin-ΔCH and βII-spectrin-ΔPH could not (Fig. 2h-k), indicating that both the actin-binding and membrane-binding functions of βII-spectrin are required for the MPS formation and/or maintenance.

### Role of the MPS in axon diameter regulation through interactions with the non-muscle myosin II (NMII)

Another family of actin-binding proteins identified by our co-IP and mass spectrometry experiments as candidate MPS-interacting proteins were NMII motor proteins, which were expressed throughout the axons of cultured neurons (Supplementary Fig. 5a). Phosphorylated myosin regulatory light chain 2, an activator of contractile NMII, has been shown to colocalize with actin rings in the AIS of cultured neurons^28, 29^, suggesting that the NMII bipolar filaments could bind to actin rings in the MPS and potentially generate tension to control the morphology and contractility of neurites. The NMII bipolar filaments could interact with MPS in two potential binding modes: the motor domains at the two ends of the 300-nm long NMII bipolar filaments^30-34^ may bind to the same actin ring or connect different actin rings in the MPS, providing the possibilities to control the radial and longitudinal contractility of axons, respectively. It remains unclear which of these two is the major binding mode.

To probe how NMII binds to the MPS, we used two-color STORM to examine the center and ends of the NMII bipolar filaments (marked by the C- and N-terminus of NMII heavy chain, respectively) with respect to the MPS (Fig. 3a). If each NMII filament binds within the same actin ring in the MPS (the first binding mode), both C- and N-terminal labels of NMII would be colocalized with the actin rings and hence form alternating patterns with the spectrin tetramer centers, marked by the C-terminus of βII-spectrin (Fig. 3a).

**Fig. 3:**
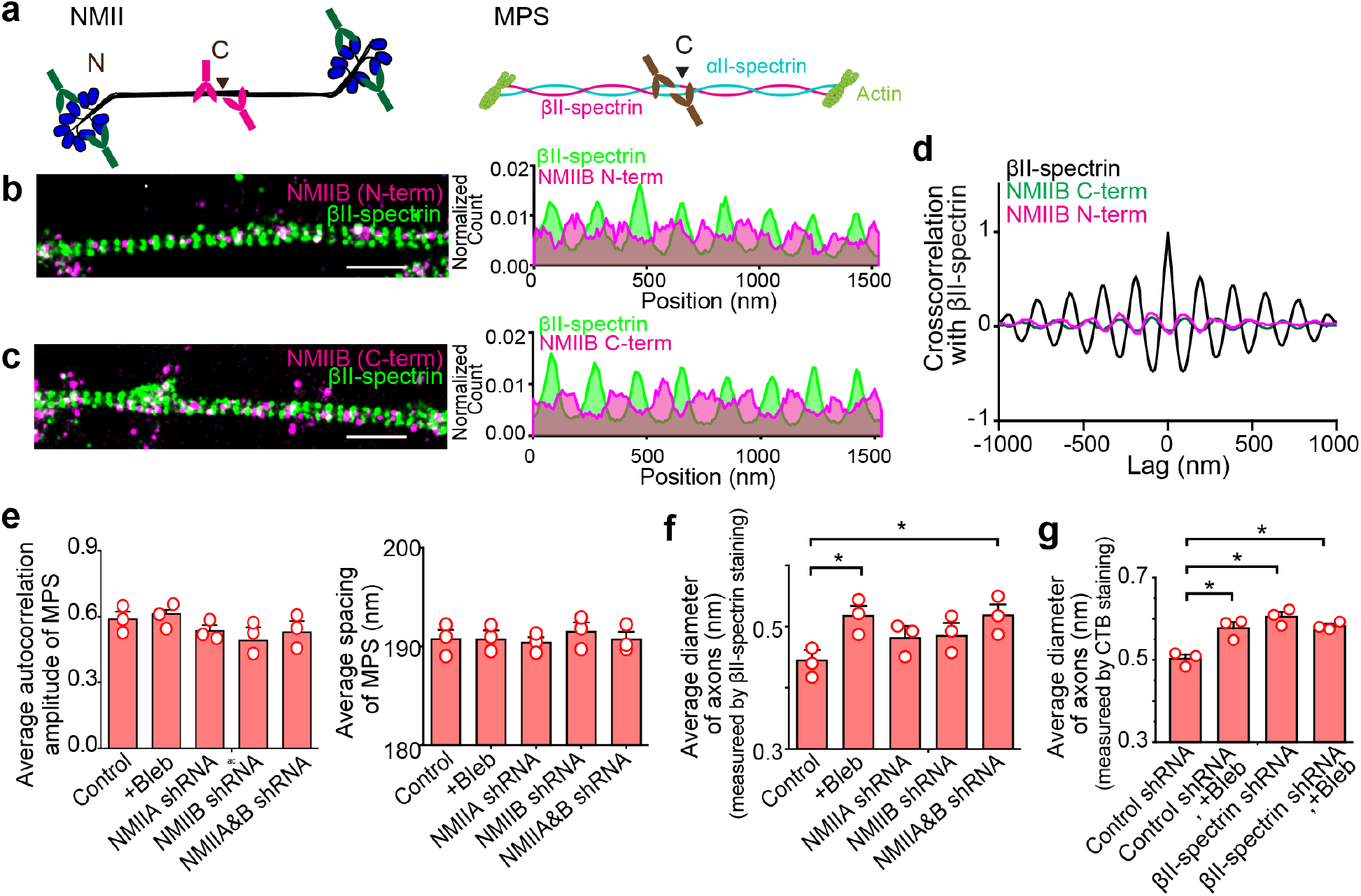
Non-muscle myosin II (NMII) filaments binds to the MPS and plays a role in radial contractility of axons. **a** Diagrams showing the antibody-binding epitopes at the C- or N-terminus of a NMII bipolar filament (left), and the antibody-binding epitope at the C-terminus of βII-spectrin located near the center of the spectrin tetramer. **b** Left: Two-color STORM image of βII-spectrin (C-terminus, green) and NMIIB (N-terminus, magenta) in axons of cultured hippocampal neurons. Right: Average 1D distributions of βII-spectrin (C-terminus, green) and NMIIB (N-terminus, magenta) signals projected to the longitudinal axon axis from many axon segments. The axonal regions were aligned based on the phase of the βII-spectrin distribution for each analyzed region and the average 1D distributions were then calculated for both βII-spectrin and NMIIB. Scale bar: 1 μm. **c** Similar to (**b**) but the neurons were immunostained with the antibody that binds to the C-terminus instead of N-terminus of NMIIB. **d** Average 1D cross-correlation functions between the distributions of βII-spectrin (C-terminus) and NMIIB (N-terminus) (magenta) and between the distributions of βII-spectrin (C-terminus) and NMIIB (C-terminus) (green) along the axons of cultured neurons, derived from many axon segments. The average 1D auto-correlation function of βII-spectrin distribution (black) is shown as a reference. **e** Average 1D autocorrelation amplitudes of βII-spectrin distribution (left) and average periodic spacing of the MPS (right), for the axons of untreated neurons, blebbistatin (Bleb)-treated neurons, and neurons transfected with adenoviruses expressing shRNAs against NMIIA heavy chain, NMIIB heavy chain, or both NMIIA and NMIIB heavy chains. **f** Average diameter of axons (measured by βII spectrin immunostainining) for untreated neurons, Bleb-treated neurons, and neurons treated with shRNAs against NMIIA heavy chain, NMIIB heavy chain, or both NMIIA and NMIIB heavy chains. **g** Average diameter of axons (measured by cholera toxin B (CTB) staining) for neurons treated with control (scramble) shRNA, neurons treated with control shRNA and Bleb, neurons treated with βII-spectrin shRNA, and neurons treated with βII-spectrin shRNA and Bleb. Data are mean ± s.e.m (*n* = 3 biological replicates; 50-100 axonal regions were examined per condition). * indicates *p* < 0.05 and ** indicates *p* < 0.005 (unpaired student’s t-test).

For the second binding mode, if each NMII filament connects adjacent actin rings, only the N-terminus of NMII would be colocalized with the actin rings but the C-terminus of NMII would be colocalized with the spectrin tetramer centers. Our STORM images showed that both C- and N-termini of NMIIB, and the N-terminus of NMIIA (a good antibody against C-terminus of NMIIA is lacking), preferentially localized midway between the periodic stripes formed by the spectrin tetramer centers, which is where the actin rings are situated (Fig. 3b, c and Supplementary Fig. 5b). The average cross-correlation function between the 1D distributions of βII-spectrin (C-terminus) and NMII (N-terminus or C terminus) signals along the longitudinal axon axis reached minimum at the spectrin tetramer centers and maximum at midway between adjacent spectrin tetramer centers (Fig. 3d and Supplementary Fig. 5b). These results suggest that among the two binding modes, the first mode where the NMII filaments bind within the same actin rings is the preferred one. The sparse appearance of NMII filaments in the STORM images could be because the labeling efficiency of the NMII antibodies is relatively low, or the number of NMII filaments bound to each actin ring is small. It is also worth noting that there is a considerable non-zero baseline in the average 1D distribution of NMII (N-terminus or C terminus) along the axon axis (Fig. 3b, c and Supplementary Fig. 5b). This non-zero baseline could represent 1) NMII molecules randomly distributed in axons (i.e., unbound NMII or NMI filaments bound to non-MPS actin filaments), 2) nonspecific binding of these NMII antibodies, and/or 3) a fraction of NMII filaments adapting the second binding mode where the NMII bipolar filaments connect adjacent actin rings in the MPS. In addition, in the second binding mode, if the NMII filaments could bridge every other actin ring spaced ∼380 nm apart, both C- and N-terminal labels of NMII would also appear to be colocalized with the actin rings. Although we cannot rule out this possibility, we consider this scenario less likely because it would require all NMII filaments connecting every other actin ring to be stretched to ∼380 nm long whereas the mean length of NMII bipolar filaments was measured to be 300 nm with a standard deviation of 20 nm by super-resolution fluorescence imaging^34^, or 290-320 nm with a standard deviation of 20-30 nm by electron microscopy^30, 33^.

Next, we examined whether the axon diameter is changed upon inhibition of the contraction activity of NMII by treatment with Blebbistatin (Bleb), a drug that prevents the transition of NMII filaments into tension-generating states without necessarily dissociating NMII from actin^35^. Bleb treatment (Supplementary Fig. 5c) did not significantly disrupt the MPS structure or its 190-nm spacing (Fig. 3e and Supplementary Fig. 5d), but increased the average axon diameter by 20-30% (Fig. 3f). In addition to Bleb treatment, knockdown of both NMIIA and NMIIB (Supplementary Fig. 5e) also led to a similar increase in the average axon diameter without changing the average spacing between adjacent actin rings (Figs. 3e, 3f and Supplementary Fig. 5d). These results are consistent with the model that NMII filaments bind to actin filaments within individual rings, forming actomyosin complexes that radially contract the actin rings and control the axon diameter. In parallel to our work, two independent studies were reported recently showing that NMII filaments interact with the MPS and regulate the diameter of axons^36, 37^.

To further assess whether the effect of NMII inhibition is through the interaction of NMII with the MPS or with other actin structures, we further examined whether Bleb treatment could also increase the average axon diameter of neurons in which the MPS had been disrupted by βII-spectrin knockdown (Supplementary Fig. 5f). In the experiments described above, we used βII-spectrin immunostaining to estimate the axon diameter. Here, to allow measurements of the axon diameter in βII-spectrin knockdown neurons, we used cholera toxin B (CTB) to label the axonal membrane. We found that βII-spectrin knockdown led to an 20-30% increase in the average axon diameter (Fig. 3g and Supplementary Fig. 5g) without significantly changing the expression levels of NMIIA and NMIIB in axons (Supplementary Fig. 5e). The degree of axon diameter enlargement induced by βII-spectrin knockdown was similar to that induced by Bleb treatment or by NMIIA and NMIIB knockdown of WT neurons. Furthermore, Bleb treatment did not further increase the average axon diameter of βII-spectrin knockdown neurons (Fig. 3g), suggesting a lack of MPS-independent mechanisms for the NMII-mediated axon contraction.

### Transmembrane proteins and membrane-associated signaling proteins interacting with the MPS

Among the 515 candidate MPS-interacting proteins commonly identified from the four pulldown experiments (three with mouse brain tissues using αII-spectrin, βII-spectrin, or α-adducin as the bait, and one with cultured neurons using βII-spectrin as the bait) by mass spectrometry, only 23 were transmembrane proteins and some transmembrane proteins known to be associated with the MPS, such as sodium channels^1^, were missing. The low abundance of transmembrane proteins identified was likely because transmembrane proteins are difficult to immunoprecipitate due to their low solubility in the co-IP lysis buffer. To obtain a more inclusive list of candidate MPS-interacting transmembrane proteins, we pooled together all transmembrane proteins identified from the four pulldown experiments, rather than only selecting the common proteins detected in all four pulldown experiments. The resulting pooled list contained 186 transmembrane proteins, among which several have been previously shown to bind to the MPS and exhibit periodic distributions at the AIS or the node of Ranvier, including the sodium channel Na_v_, potassium channel K_v_1.2, and two cell adhesion molecules (neurofascin and NrCAM)^1, 6, 11, 12^. GO term enrichment analyses of these 186 transmembrane proteins showed enrichment for ion channel activity, glutamate and GABA receptor activity, membrane transporter and symporter activity, G-protein coupled receptor (GPCR) activity, cell-cell adhesion activity, and receptor protein kinase activity (Fig. 4a).

**Fig. 4:**
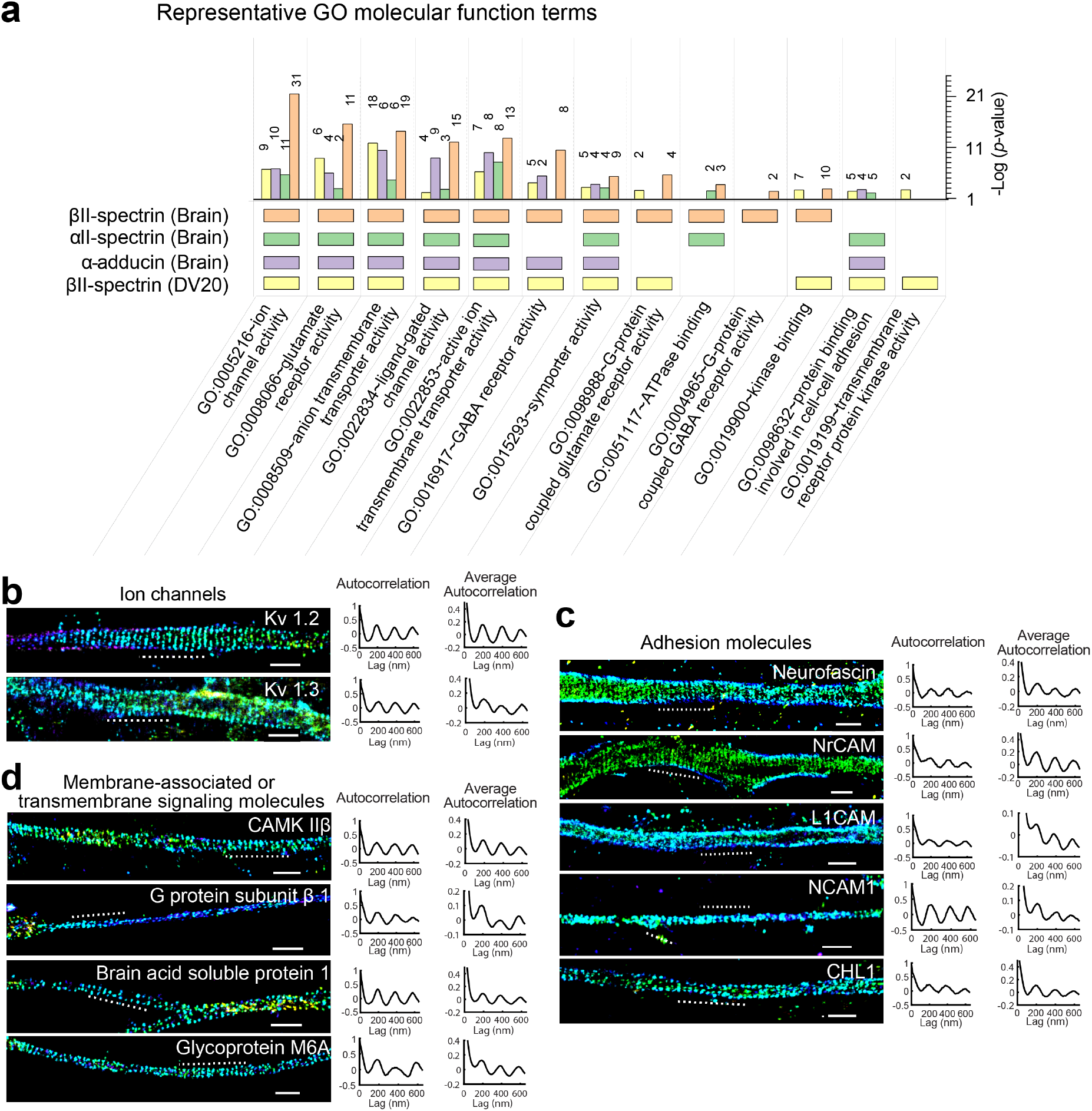
Proteomic analysis and validation of MPS-interacting transmembrane or signaling proteins. **a** Enriched GO molecular function (MF) terms of mass-spectrometry-identified transmembrane proteins in the four co-IP experiments, including one from cultured hippocampal neurons (DIV 20) using βII-spectrin as the bait (yellow), and three from the mouse brain using αII-spectrin (green), βII-spectrin (orange), and α-adducin (purple) as the bait. Enriched GO MF terms were generated using the DAVID 6.8 platform and representative GO MF terms are shown, together with the *p*-value (bar plots on the right) and gene count (numbers next to the bars) for each GO term. **b** Left: 3D STORM images of two potassium channel subunits, K_v_ 1.2 and K_v_ 1.3, in axons. Middle: 1D autocorrelation functions of the imaged molecules for the axon region indicated by the dashed line in the left panels. Right: Average 1D autocorrelation functions of the imaged molecules over many randomly chosen axon regions. **c** Same as (**b**) but for five cell adhesion molecules, including neurofascin, NrCAM, L1CAM, NCAM1 and CHL1. **d** Same as (**b**) but for three membrane-associated (non-transmembrane) signaling molecules, including calcium/calmodulin-dependent protein kinase type IIβ (CAMK IIβ), heterotrimeric G protein β-subunit 1, and brain acid soluble protein 1, as well as a transmembrane signaling molecule, glycoprotein M6A. K_v_ 1.2, neurofascin, NrCAM, CHL1 were immunolabeled with their corresponding antibodies, and the other proteins were labeled using moderate expression of GFP-tagged proteins through low-titer lentiviral transfection of WT neurons, which were in turn immunolabeled with anti-GFP antibody. Scale bars: 1 µm.

Although the pooling described above can generate a more inclusive list of transmembrane proteins, it may also cause a concomitant rise in false positives. To validate interactions of some of these transmembrane proteins with the MPS, we applied STORM imaging to visualize several representative candidate proteins with different known cellular functions. In the ion channel category, we observed that K_v_1.2 and K_v_1.3, exhibited periodic distributions with 190-nm spacing along axons (Fig. 4b). In the cell adhesion molecule category, we observed that neurofascin, NrCAM, L1CAM, NCAM1 and CHL1 exhibited periodic distributions with 190-nm spacing and different degrees of periodicity as indicated by the autocorrelation amplitude (Fig. 4c). Compared to neurofascin, NrCAM and CHL1, the relatively low degree of periodicity for L1CAM and NCAM1 could be due to the challenges in labeling as described earlier (Supplementary Fig. 3). Unlike neurofascin, NrCAM and CHL1, which were labeled using antibodies against the endogenous proteins, L1CAM and NCAM1 were labeled using expression of GFP-fusion proteins through low-titer lentiviral transfection followed by immunolabeling using anti-GFP antibody because we could not identify antibodies against L1CAM and NCAM1 with sufficient specificity. Alternatively, the relatively low degree of periodicity could be also due to the existence of a considerable fraction of the examined molecules that were not localized to the MPS. As we have shown previously, NCAM1 shows relatively weak colocalization with the MPS under unstimulated conditions, and the binding of antibody to the extracellular domain of NCAM1, mimicking homophilic or heterophilic binding of NCAM1 during the establishment of neurite-neurite contacts, causes a substantial increase in the degree of colocalization between NCAM1 and the MPS^13^.

Since GO term analyses revealed that some candidate MPS-interacting proteins are involved in cell signaling, we also imaged several non-transmembrane, but membrane-associated signaling proteins from the candidate MPS-interacting protein list. Interestingly, we found that calcium/calmodulin-dependent protein kinase IIβ (CAMK IIβ) exhibited periodic distributions with 190-nm spacing along axons (Fig. 4d). The heterotrimeric G protein subunits also exhibited a tendency, albeit weak, to adopt the 190-nm periodic distributions (Fig. 4d). In addition, a membrane-associated signaling protein important for axon outgrowth, brain acid soluble protein 1 (BASP1)^38^, as well as a transmembrane axon-growth-related protein, glycoprotein M6A^39^, also exhibited periodic distributions along axons (Fig. 4d). These results suggest that the MPS may play a role in various signaling pathways in neurons. Indeed, we recently found that the MPS can function as a signaling platform to recruit various signaling molecules, including GPCRs, cell adhesion molecules, and RTKs, in response to the extracellular stimuli, enabling the downstream extracellular-signal-regulated kinase (ERK) signaling^13^. Results here suggest the possibility that the MPS may also play a similar role in regulating the signaling pathways involving CAMK IIβ, BASP1 and glycoprotein M6A.

### Role of the MPS in neurite-neurite interactions mediated by cell adhesion molecules

In the brain, cell adhesion molecules play essential roles in regulating neuronal migration and neurite outgrowth, and in establishing and maintaining various types of neuron-neuron contacts, such as axon-axon fasciculation (i.e. bundling), axon-dendrite fasciculation, as well as synaptic interactions^40-44^. Because our experiments revealed MPS interactions with several cell adhesion molecules, it raises the possibility that the MPS serves as a structural platform to anchor these molecules and facilitate adhesion-molecule-mediated contacts between neurites.

If the MPS-associated adhesion molecules indeed can bring neurites into contact, we expect that the MPS structures in the abutting neurites would be aligned in phase with each other by the contacting pairs of adhesion molecules, which bind to defined sites in the MPS. Indeed, we observed that the periodic distributions of βII-spectrin in the abutting axon-axon and axon-dendrite pairs tended to be in phase (Supplementary Fig. 6), consistent with the previous observation of in-phase alignment of the MPS of two contacting axons^12^.

To test whether the MPS structure is indeed involved in the formation of axon-axon and axon-dendrite interactions, we examined how axon-axon and axon-dendrite fasciculations were affected by βII-spectrin knockdown, which is known to disrupt the MPS^4, 7^, or by knocking down ankyrin B, a protein that can interact with both the MPS^1^ and cell adhesion molecules^2^ and hence may serve as an adaptor to anchor these adhesion molecules to the MPS.

Notably, axon-axon bundling was greatly reduced by either βII-spectrin or ankyrin B knockdown (Fig. 5a). In order to quantify the axon-bundling effect, we used an automated algorithm to determine the average width of the axon-positive areas in the images (including both axon bundles and single axons), and found that this width parameter was significantly decreased in both βII-spectrin knockdown neurons and ankyrin B knockdown neurons, as compared to control neurons treated by scrambled shRNA (Fig. 5a). The average width values observed in these βII-spectrin knockdown and ankyrin B knockdown samples reduced to values that were close to those observed from single, unbundled axons (Fig. 5a), indicating a substantial disruption of axon-axon bundling when the MPS was disrupted or when the adaptor protein between cell adhesion molecules and the MPS was depleted. To examine the knockdown effects on axon-dendrite interactions, we quantified two properties: 1) axon-dendrite bundling, defined by the average fraction of dendrite segments that had adhering axons; and 2) the density of synaptic contacts along dendrites. We imaged synaptic contacts using immunofluorescence against presynaptic scaffolding protein bassoon and postsynaptic density protein homer1 and counted puncta that were positive of both bassoon and homer1 as synapses. We found that βII-spectrin knockdown and ankyrin B knockdown caused a significant decrease both in the degrees of axon-dendrite bundling and in the synapse density along dendrites (Fig. 5b, c). These data suggest that the MPS plays a role in axon-axon and axon-dendrite interactions.

**Fig. 5:**
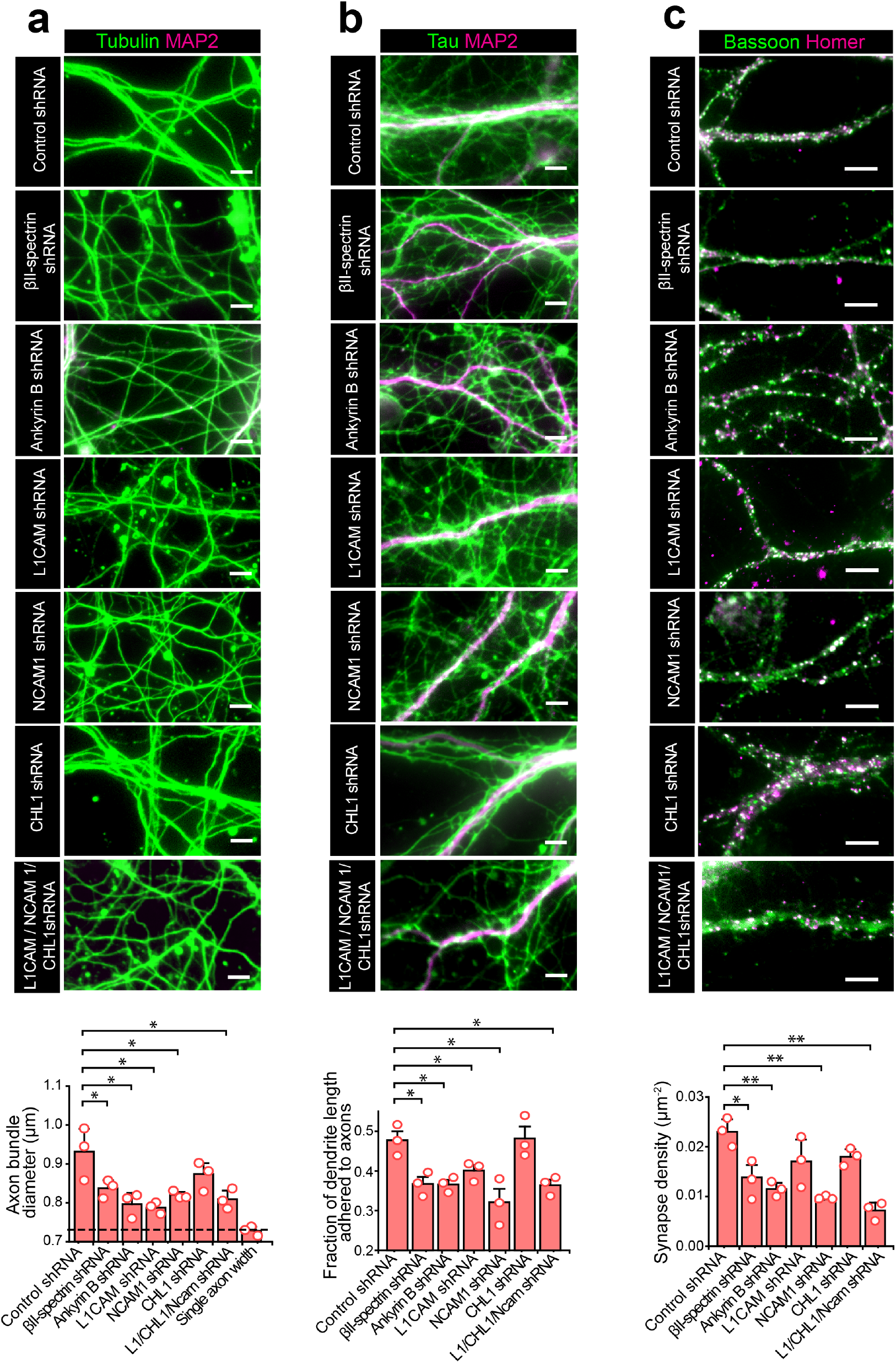
The MPS plays a role in axon-axon interactions, axon-dendrite interactions and synapse formation. **a** Top panels: Conventional fluorescence images of tubulin (green) and the dendrite marker MAP2 (magenta) for neurons transfected with adenoviruses expressing scrambled (control) shRNA, βII-spectrin shRNA, ankyrin B shRNA, L1CAM shRNA, NCAM1 shRNA, or CHL1 shRNA, as well as neurons transfected with adenoviruses expressing shRNAs against all three adhesion molecules (L1CAM, NCAM1 and CHL1). The tubulin-positive neuronal processes lacking MAP2 signal are axons. Bottom panel: Average diameter of the axon bundles (or single axons) quantified for the seven conditions described in the top panels. The last bar and dashed line represent the average single-axon diameter obtained from the fluorescence image of sparsely cultured neurons. **b** Top: Conventional fluorescence images of cultured neurons immunostained for the axon marker Tau (green) and the dendrite marker MAP2 (magenta) under the seven conditions described in (**a**). Bottom: Average fraction of the total length of dendrites that are bundled with axons, quantified for the seven conditions indicated in the top panel. **c** Top: Conventional fluorescence images of cultured neurons immunostained for the presynaptic marker Bassoon (green) and the postsynaptic marker Homer1 (magenta) under the seven conditions described in (**a**). Bottom: Average synapse density per unit area on dendrites, quantified for the seven conditions as described in the top panel. Only puncta that show both presynaptic and postsynaptic marker signals were counted as synapses. Data are mean ± s.e.m (*n* = 3 biological replicates; 10-20 imaged regions were examined per condition). * indicates *p* < 0.05 and ** indicates *p* < 0.005 (unpaired student’s t-test). Scale bars: 5 µm.

Next, we examined the effect of knocking down MPS-associated cell adhesion molecules on axon-axon and axon-dendrite interactions. Among the adhesion molecules that we have validated as interacting partners of the MPS (Fig. 4c), we examined two that are known to be enriched in axons, L1CAM and CHL1^45-47^ and one that is enriched in both axons and dendrites, NCAM1^45^, by knocking down these proteins (Supplementary Fig. 7). We did not study the effect of knocking down neurofascin and NrCAM because they are only enriched in the AIS region^6, 11^, a relatively short segment in axons. Knockdown of L1CAM or NCAM1 decreased the axon-axon and axon-dendrite bundling, whereas knockdown of CHL1 did not show any significant effect (Fig. 5a, b). The density of synapses was reduced only in the NCAM1 knockdown neurons, but not in the L1CAM or CHL1 knockdown neurons (Fig. 5c), consistent with the results from previous studies^46-48^. When all three adhesion molecules were knocked down, axon-axon bundling, axon-dendrite bundling, and synapse density were all reduced (Fig. 5a-c). These data suggest that L1CAM and NCAM1 are involved in axon-axon and axon-dendrite fasciculations, that NCAM1 is additionally involved in synapse formation, whereas CHL1 is not required for any of these three types of neurite interactions.

Our βII spectrin knockdown and ankyrin B knockdown experiments showed that the MPS plays a role in axon-axon and axon-dendrite interactions, and our L1CAM and NCAM1 knockdown experiments, as well as previous studies^42, 44, 47, 48^, implicated these cell adhesion molecules in axon-axon and axon-dendrite contacts. An interesting question arises as to whether the role of the MPS in axon-axon and axon-dendrite contacts is simply to maintain proper expression levels of these cell adhesion molecules at the neurite surface. We thus examined whether the observed effect of βII-spectrin knockdown or ankyrin B knockdown on axon-axon and axon-dendrite interactions could be simply explained by a reduction in the cell-surface expression levels of the cell adhesion molecules in these knockdown neurons by measuring the amounts of L1CAM, NCAM1 and CHL1 at the surface of neurites. βII-spectrin or ankyrin B knockdown only led to a small reduction in the cell-surface expression levels of L1CAM, but did not decrease the cell-surface expression levels of NCAM1 or CHL1 (Supplementary Fig. 8). It is thus unlikely that this small effect on the cell-surface expression levels of cell adhesion molecules could fully account for the substantial reduction in axon-axon bundling, axon-dendrite bundling, and synapse density observed in βII-spectrin knockdown or ankyrin B knockdown cells. Hence, our data suggest that spatial organization of the cell adhesion molecules by the MPS likely help establish and/or maintain efficient molecular contacts at the interface of abutting neurites.

## Discussion

In this study, we combined co-IP and mass spectrometry to identify proteins associated with the neuronal MPS at the proteomic scale. We identified hundreds of potential candidate MPS-interacting proteins, including cytoskeletal proteins and their binding proteins, transmembrane proteins, and membrane-associated signaling molecules. The GO-term analyses suggest that these candidate MPS-interacting proteins are involved in a variety of molecular functions and cellular processes, including membrane organization, vesicle-mediated transport, synaptic transmission, cell morphogenesis, differentiation and communication, cell adhesion and migration, brain development, etc. Using super-resolution microscopy, we imaged ∼20 of these candidate MPS-interacting proteins, including structural components of the MPS that can bind to actin filaments, non-muscle myosin II (NMII) motor proteins, cell-adhesion molecules, ion channels, and other signal transduction related proteins, and validated their interactions with the MPS.

Based on these results, and prior knowledge of the neuronal MPS^49^ and the erythrocyte membrane skeleton^2, 3^, we propose a more comprehensive picture of the molecular architecture of the MPS and its interacting proteins in neurons (Fig. 6). The neuronal MPS is comprised of actin rings periodically spaced by spectrin tetramers along the neurites. The actin ring consists of actin filaments, actin-capping proteins, including adducin and tropomodulins, and other actin-binding proteins, including dematin and coronins. In addition, many transmembrane proteins, such as ion channels, cell adhesion molecules, receptors, and glycoproteins, also interact with the neuronal MPS. These transmembrane proteins may be associated with either the actin rings through dematin as adaptors or associated with the spectrin tetramers through ankyrins as adaptors, and both association modes have been previously observed to connect some transmembrane proteins to the erythrocyte membrane skeleton^2, 3^.

**Fig. 6:**
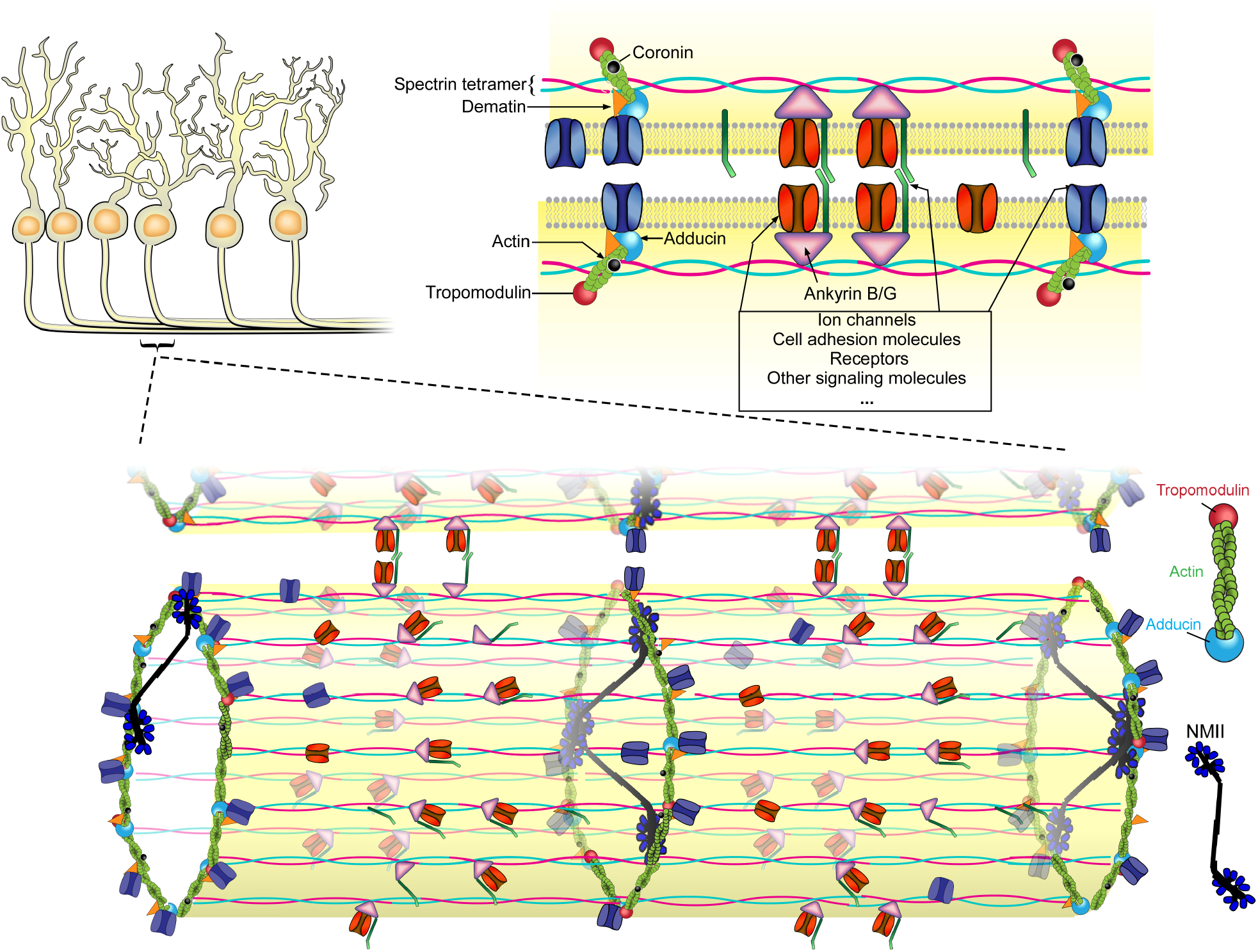
Molecular architecture model of the neuronal MPS in the neurite-neurite fasciculations. In neurites, actin filaments form ring-like structures that are connected by spectrin tetramers, and actin filaments in these rings are likely capped by adducin and tropomodulin at their fast- and slow-growing ends, respectively. The length and structure of the actin filaments within the actin rings remains an open question. Dematin and coronin bind to the actin filaments, and ankyrin B and ankyrin G bind to the site near the center of each spectrin tetramer. The NMII bipolar filaments bind preferentially to actin filaments within the actin rings and can regulate the diameter of neurites by exerting radial contractile forces. It is possible that a smaller fraction of NMII bipolar filaments may connect the two adjacent actin rings. Membrane proteins, such as ion channels, cell adhesion molecules, receptors and signaling molecules, are associated either with the actin filaments through adaptor proteins dematin, or associated with the center positions of spectrin tetramers through the adaptor protein ankyrin, to form actin and ankyrin junctional complexes at the plasma membrane of neurites. Recruitment of cell adhesion molecules to periodic sites on the MPS and interactions of cell adhesion molecules between two abutting neurites bring their respective MPS structures in phase, which could in turn bring additional MPS-bound cell adhesion molecules from the two neurites into proximity, enhancing neurite-neurite interactions. Moreover, the MPS could serve as a structural platform that recruits and organizes transmembrane proteins and membrane-associated signaling molecules, potentially facilitating a variety of signaling pathways in neurons.

Compared to the erythrocyte membrane skeleton, the neuronal MPS exhibits several notable distinctions. First, in addition to a few shared structural components (tropomodulin 1, *α*-adducin, and dematin) used by both structures, the neuronal MPS contains distinct protein homologs and unique proteins. For example, αII-spectrin, βII-spectrin, βIII-spectrin, βIV-spectrin, ankyin B, and ankyrin G are protein homologs specifically observed in the neuronal MPS, whereas the counterparts in the erythrocyte membrane skeleton are αI-spectrin, βI-spectrin and ankyrin R, respectively. Moreover, there are many unique neuronal MPS-associated proteins that have not been found on the erythrocyte membrane skeleton, such as structural proteins (e.g. coronins), transmembrane proteins (e.g. sodium and potassium ion channels, GPCRs, RTKs), and membrane-associated signaling molecules (e.g. CAMK IIβ). Ultrastructure wise, the erythrocyte membrane skeleton adopts a two-dimensional polygonal lattice structure^2, 3^, whereas the neuronal MPS adopts a one-dimensional periodic structure in axons and portions of dendrites and a two-dimensional polygonal lattice structure in the neuronal cell body and portions of dendrites^1, 4, 6, 7^. We observed actin-capping proteins in the one-dimensional periodic MPS structure in neurites, including both adducin and tropomodulin. It has been shown previously that adducin binds to the fast-growing-end of actin filaments 25-fold stronger than to the sides of actin filaments^50^, and tropomodulin, in conjunction with tropomyosin, is a slow-growing end capping protein and cannot bind along the sides of actin filaments^51^. Our observations thus suggest the possibility that the actin filaments in the periodic actin rings in the neuronal MPS are capped at both ends. However, it remains an open question whether the lengths of these actin filaments in neuronal MPS are short 12-17mers as observed for the erythrocyte membrane skeleton^3^, or whether their lengths could be longer and more heterogeneous^29^.

Moreover, we observed that tension-generating non-muscle myosin II (NMII) motors are associated with the neuronal MPS. Our observation that both C- and N-terminus of the NMII preferentially colocalized with the actin rings favors the model that the NMII biopolar filaments bind within individual actin rings, although our data do not exclude the possibility that a smaller fraction of NMII biopolar filaments connect the adjacent actin rings in the MPS (Fig. 6). Therefore, NMII filaments potentially can form actomyosin complex with the actin filaments in individual actin rings and exert contractile forces to control the diameter of the neurite. Indeed, we observed that acute inhibition of NMII activity by Blebbistatin treatment, depletion of both NMIIA and NMIIB heavy chains, or depletion of βII-spectrin all resulted in 20-30% increase of average axon diameter. Acute inhibition of NMII activity by Blebbistatin did not increase the axon diameter of βII-spectrin knockdown neurons, suggesting that the NMII-mediated radial contractility of axons is unlikely through an MPS-independent mechanism. Consistent with our observations, two parallel studies reported recently showed that NMII filaments interact with the MPS and that inhibition of NMII increases the average axon diameter^36, 37^, although the data presented in these two studies cannot unambiguously point to which of the two binding modes is the major binding mode of NMII bipolar filaments to the MPS structures^36, 37^. Together, our results and these parallel studies showed that the interaction of NMII filaments with the MPS could control radial contractility of axons. This radial contractility of axons could be functionally important, because the action potential conduction velocity along axons can change substantially upon a 10-20% axonal diameter change^52, 53^. Indeed, inhibiting NMII activity increases axonal signal propagation velocity^37^. In addition, this NMII-mediated axon contractility has also been shown to affect cargo transport in axons^36^. Moreover, because axonal tension and mechanosensing are critical for axonal growth and synaptic plasticity^54, 55^, this NMII-MPS interaction may also be important for these processes.

Finally, we showed that the MPS plays a role in the establishment or maintenance of axon-axon and axon-dendrite interactions. MPS organizes cell adhesion molecules, such as L1CAM and NCAM1, into periodic distributions along neurites, and knockdown of L1CAM or NCAM1 led to a substantial reduction in axon-axon and axon-dendrite bundling. Importantly, disruption of the MPS by βII-spectrin knockdown or depletion of ankyrin B, the adaptor protein which connects cell adhesion molecules to the MPS, also caused a substantial reduction in axon-axon and axon-dendrite bundling, as well as in synaptic contacts between axons and dendrites, even though the cell-surface expression levels of these cell-adhesion molecules did not decrease substantially under these conditions. These results suggest that the mechanism by which the MPS facilitates axon-axon and axon-dendrite interactions goes beyond maintaining the proper amount of cell adhesion molecules at the surface of neurites. We propose the following potential mechanisms for the MPS-dependent axon-axon and axon-dendrite interactions. First, recruitment of adhesion molecules to periodic sites on the MPS may substantially enhance the interactions of adhesion molecules between two abutting neurites, because contacts between some MPS-bound adhesion molecules from two abutting neurites may lead to in-phase alignment of the MPS structures contained in the two neurites, which may in turn bring many additional MPS-bound adhesion molecules from the two neurites into proximity. In addition, it has been shown that downstream signaling pathways activated by adhesion molecules are required for axon outgrowth, guidance and fasciculation^43, 56, 57^. It is thus also possible that MPS facilitates the neurite-neurite interactions by providing a structural platform to recruit relevant signaling molecules and adhesion molecules to common sites to form signaling complexes and enable signaling relevant to neurite-neurite interactions. In support of this notion, we have recently shown that the MPS can function as a platform to recruit NCAM1 and related signaling molecules to enable NCAM1-induced ERK signaling^13^. Our mass spectrometry and super-resolution imaging results here further showed that many other signaling molecules are associated with the MPS, suggesting the possibility that the MPS could facilitate other signaling pathways involved in the establishment of neurite-neurite interactions. Together, our results provide new insights into how axon-axon and axon-dendrite interactions are mediated at the molecular level. More generally, our observations that many transmembrane proteins and membrane-associated signaling molecules are organized by the MPS suggests that the MPS could serve as a structural platform to facilitate a variety of signaling pathways in neurons.

## Methods

### Primary culture of mouse hippocampal neurons

All experimental procedures were performed in accordance with the Guide for the Care and Use of Laboratory Animals of the National Institutes of Health. The protocol was approved by the Institutional Animal Care and Use Committee (IACUC) of Harvard University.

Primary cultures of hippocampal neurons were prepared as previously described ^10^. The mouse lines used in this study include CFW mice (Strain code 024, Charles River Laboratories), a *βII-Spec*^*flox/flox*^ mouse line^27^, a Nestin-Cre mouse line (Stock number 003771, The Jackson Laboratory), a whole body Tmod1 knockout mouse line that expresses a *Tmod1* transgene only in the heart^58^, a whole body *Tmod2* knockout mouse line^59^, and a whole body *Dmtn* knockout mouse line^60^. Hippocampi were isolated from mouse embryos (E18) or newborn pups (P0) and digested with 0.25% trypsin-EDTA (1x) (Sigma, T4549) at 37^°^C for 15 minutes. The digested tissues were washed in Hanks’ Balanced Salt Solution (HBSS) (Thermo Fisher Scientific, 14175079) for three times, and then transferred to culture medium consisting of Neurobasal medium (Thermo Fisher Scientific, 21103049) supplemented with 37.5 mM NaCl, 2% (vol/vol) B27 supplement (Thermo Fisher Scientific, 17504044) and 1% (vol/vol) Glutamax (Thermo Fisher Scientific, 35050-061). The tissues were gently triturated in the culture medium until there were no chunks of tissue left. Dissociated cells were then counted and plated onto poly-D-lysine-coated 18-mm coverslips or 10 cm petri dishes. Cultures were maintained in the culture medium in a humidified atmosphere of 5% CO_2_ at 37°C. The neurons were fed with one-half medium volume change every five days.

### U2OS cell culture

Human bone osteosarcoma epithelial cells (U2OS cells) were obtained from ATCC. U2OS cells were plated onto 8-well Lab-Tek glass bottom chambers (Thermo Fisher Scientific, 155409) and maintained in complete growth Dulbecco’s modified Eagle’s medium (DMEM), supplemented with 10% (vol/vol) fetal bovine serum (FBS), 0.1% (vol/vol) penicillin–streptomycin.

### Antibodies

The following primary antibodies were used in this study: guinea pig anti-MAP2 antibody (Synaptic Systems, 188004), rabbit anti-MAP2 antibody (Synaptic Systems, 188002), mouse anti-αII spectrin antibody (Biolegend, 803201), mouse anti-βII spectrin antibody (BD Biosciences, 612563), mouse anti-dematin antibody (Santa Cruz Biotechnology, sc-135881), rabbit anti-coronin 2B antibody (Novus Biologicals, NBP 1-85567), mouse anti-tubulin antibody (Santa Cruz Biotechnology, sc-5286), rabbit anti-Tau antibody (Synaptic Systems, 314002), mouse anti-K_v_1.2 channel antibody (Neuromab, 75-008), rabbit anti-neurofascin antibody (Neuromab, 75-172), rabbit anti-NrCAM (Abcam, ab24344), goat anti-CHL1 antibody (R&D systems, AF2147), rabbit anti-NCAM1 antibody (EMD Millipore, AB5032), mouse anti-ankyrin G antibody (Santa Cruz Biotechnology, sc-12719), mouse anti-bassoon antibody (Enzo, ADI-VAM-PS003-F), rabbit anti-homer antibody (Synaptic Systems, 160003), rabbit anti-protein 4.1B antibody (LifeSpan BioSciences, LS-C83790-100), rabbit anti-L1CAM antibody (ABclonal, A8555), rat anti-L1CAM antibody (R&D Systems, MAB5674), rabbit anti-Myh10 (N-terminus) antibody (GeneTex, GTX133378), abbit anti-Myh9 (N-terminus) antibody (GeneTex, GTX101751), rabbit anti-Myh10 (C-terminus) antibody (Biolegend, 909901), rabbit anti-GFP antibody (Thermo Fisher Scientific, A11122). rabbit anti-β-actin antibody (Proteintech, 20536-1-AP).

The following secondary antibodies were used in this study: CF680-conjugated donkey anti-mouse IgG antibody (Biotium, 20819), Alexa-647-conjugated donkey anti-mouse IgG antibody (Jackson ImmunoResearch, 715-605-151), Alexa-647-conjugated donkey anti-rabbit IgG antibody (Jackson ImmunoResearch, 711-605-152), Alexa-647-conjugated donkey anti-goat IgG antibody (Jackson ImmunoResearch 705-605-147), Alexa-647-conjugated donkey anti-rat IgG antibody (Jackson ImmunoResearch, 712-605-153), Cy3-conjugated donkey anti-rabbit IgG antibody (Jackson ImmunoResearch, 711-165-152), Cy3-conjugated donkey anti-guinea pig IgG antibody (Jackson ImmunoResearch, 706-165-148), Cy3-conjugated donkey anti-mouse IgG antibody (Jackson ImmunoResearch, 711-165-151). Alexa-488-conjugated donkey anti-guinea pig IgG antibody (Jackson ImmunoResearch, 706-545-148).

### Co-immunoprecipitation of MPS-interacting proteins for mass spectrometry analysis

The magnetic protein-G-coated beads (Thermo Fisher Scientific, 10007D) were incubated with the irrelevant (control) antibody or antibody against βII-spectrin, αII-spectrin, or α-adducin at 4 °C overnight, followed by 3x washes using the washing buffer (PBS (pH 7.4) containing 0.02% (vol/vol) Tween-20). For mouse brain co-immunoprecipitation experiments, adult mouse brains were isolated and dounce-homogenized in the co-immunoprecipitation lysis buffer containing 25 mM Tris-HCl (pH 7.4), 150 mM NaCl, 1 mM EDTA, 5 mM EGTA, 1% NP-40, 5% glycerol, 1x protease inhibitor cocktail (Thermo Fisher Scientific, 87785) on ice. For co-immunoprecipitation experiments from cultured hippocampal neurons, DIV 20 neurons were lysed on ice using the co-immunoprecipitation lysis buffer. After incubating 15 min, cell lysates were centrifuged at 13,000 g for 5 min at 4 °C. The supernatant was collected and then incubated with the antibody-coated magnetic beads for 3 hours at 4 °C. The magnetic beads with the co-immunoprecipitated products were washed 5 times using the washing buffer. Finally, the co-immunoprecipitated proteins were eluted using Laemmili sample buffer (Bio-Rad, #161-0737) for 10 min at 90 °C. The eluted protein mixtures from mouse brains or DIV20 neurons were then separated using SDS-PAGE, and each gel lane was cut into gel pieces and subjected to in-gel tryptic digestion as previously described ^61, 62^.

### Sample preparation for quantitative mass spectrometry

At DIV 20, cultured hippocampal neurons treated with adenoviruses expressing either βII-spectrin shRNA or scrambled (control) shRNA were lysed on ice using the RIPA buffer (Thermo Fisher Scientific, 89900) complemented with 1x protease and phosphatase inhibitor cocktail (Thermo Fisher Scientific, 78440). Cell lysates were transferred to microcentrifuge tubes and centrifuged at 13,000 g for 5 min at 4 °C, and the supernatants were collected for quantitative mass-spectrometry analyses. Protein mixtures (three replicates for each condition) were digested and labeled by tandem mass tags (TMT) 6plex reagent (Thermo Fisher Scientific, 90061), according to the manufacturer’s protocol.

### Protein identification using mass spectrometry

Experiments were performed on a hybrid ion trap-Orbitrap mass spectrometer (Orbitrap Velos, Thermo Fisher Scientific) for all analytical runs. HPLC samples (Waters) were injected into the trap column (75 µm column ID, 5 cm packed with 5 µm beads on 200 Å poros, from Michrom Bioresources, Inc.) and washed for 15 min, and were then eluted to the analytical column with a gradient from 2 to 32% of 0.1% formic acid over 90 min. The instrument was set up to run the TOP 20 method for MS/MS in iontrap. The raw mass spectrometry data were analyzed by Proteome Discoverer 2.1 (Thermo Fisher Scientific), and searches were performed against mouse (*Mus musculus*) Uniprot database and known common lab contaminants. 1% false discovery rate (FDR) was held for both protein and peptide levels for all output reported data.

### Plasmids construction and transfection of βII spectrin truncation mutants

βII-spectrin-ΔCH-GFP and βII-spectrin-ΔPH-GFP plasmids were modified from FUGW-GFP plasmid (Addgene, 14883, Cambridge, MA). In details, for βII-spectrin-ΔCH-GFP construct, 1-303 amino acids truncated βII-spectrin was amplified from wide-type βII-spectrin and inserted into FUGW-GFP. A GGGGS peptide linker was inserted between truncated βII-spectrin and GFP. Likewise, for βII-spectrin-ΔPH-GFP, 2169-2364 amino acids truncated βII spectrin was amplified from wide-type βII-spectrin and inserted into FUGW-GFP with a GGGGS peptide linker. Plasmids were transfected into cultured neurons at DIV 7-9 using a calcium phosphate transfection kit (Thermo Fisher, K2780-01). Experiments were performed 2 or 3 days post transfection.

### Expression of GFP fusion proteins in neurons for imaging MPS-interacting proteins

Plasmids used in this study were purchased from GeneCopoeia or OriGene, and cloned into lentiviral expression vector FUGW (Addgene, 14883), or pLVX (Clontech, 632155), through Gibson assembly reaction (New England Biolabs, E2611S) according to the manufacturer’s protocol. *Kcna3, Gnb1* and *Camk2b* were tagged at their N-terminus with the GFP gene and cloned into pLVX vector. *Add1, Tmod1, Tmod2, Epb41l3, L1cam, Ncam1, slc2a3, Gpm6a, Basp1* were tagged at their C-terminus with the GFP gene and cloned into FUGW vector.

Lentiviruses were then produced by co-transfecting HEK293T cells with 6 µg lentiviral expression vector (FUGW or pLVX), Δ8.9 vector ^63^ (gift from Prof. David Baltimore, California Institute of Technology), and 3 µg VSVG packaging vector (Addgene, 8454) in a 100 mm dish using a calcium phosphate transfection kit (CalPhos Mammalian Transfection kit, Clontech, 631312). Two days post-transfection, the supernatant was harvested, centrifuged at 3000 rpm for 10 minutes, and then concentrated with Lenti-X concentrator (Clontech, 631231), before being snap frozen in liquid nitrogen. The lentivirus expressing the desired GFP-tagged protein was added to the neuronal cultures between 5-8 days *in vitro* (DIV).

### shRNA knockdown and blebbistatin treatment

The sense sequences of the βII-spectrin shRNA are 5′-GCATGTCACGATGTTACAA-3′ and 5′-GGATGAAATGAAGGTGCTA-3′.

The sense sequence of ankyrin B is 5′-GACAAGCAGAAGTTGTCAA-3′.

The sense sequence of α-adducin is 5’-GTGACTGCATCCAGTTTGG-3’.

The sense sequence of αII-spectrin is 5’-AGCATGATGTTCAAACACT −3’.

The sense sequence of coronin 2B is 5’-CCATCACCAAGAATGTACAT −3’.

The sense sequence of protein 4.1B is 5’-CGTGACCGGCTTCGAATAA −3’.

The sense sequences of NMIIA heavy chain shRNA and NMIIB heavy chain shRNA are 5′-TACCCTTTGAGAATCTGATAC-3′ and 5′-CTTCCAATTTACTCTGAGAA-3′, respectively.

The sense sequences of L1CAM shRNA, NCAM1 shRNA and CHL1 shRNA are 5′-TGCTAGCCAATGCCTACATTT-3′, 5′-CGTTGGAGAGTCCAAATTCTT-3′ and 5′-GCAGAAGATCAGGGTGTTT-3′, respectively.

The adenovirus expressing a scramble shRNA sequence was used as a control (Vector BioLabs, 1122). For knocking down the specific gene, adenoviruses expressing the above target sequences were added to the neuronal cultures at DIV 2-5. The knockdown efficiencies were confirmed by either immunofluorescence or western blot analysis.

For blebbistatin (Bleb) treatment, cultured neurons or the U2OS cells were treated with 100 µM Bleb for 2 hours at 37 °C.

### Cell fixation and immunostaining for fluorescence imaging

Cultured hippocampal neurons and U2OS cells were fixed using 4% (w/v) paraformaldehyde (PFA) in phosphate buffered saline (PBS) for 30 min at room temperature (RT), washed three times in PBS, and permeabilized with 0.15% (vol/vol) Triton X-100 in PBS for 10 min. Neurons were then blocked in blocking buffer containing 3% (wt/vol) bovine serum albumin (BSA) in PBS for 1 hour, and subsequently stained with primary antibodies in blocking buffer overnight at 4 °C. Neurons were washed three times with PBS and stained with fluorophore conjugated secondary antibodies in blocking buffer for 1 hour at RT. The samples were post-fixed with 4% PFA for 20 min. To measure the average axon diameter, neurons were stained with either anti-βII-spectrin antibody (BD Biosciences, 612563) or Cholera Toxin Subunit B (Thermo Fisher Scientific, C34778).

For immunostaining of the cell-surface NCAM1, CHL1 and L1CAM, neurons were fixed using 4% PFA in PBS and blocked in blocking buffer for 1 hour. The primary antibody detecting the extracellular domain of NCAM1, CHL1 or L1CAM was then added to label the surface NCAM1, CHL1 and L1CAM, respectively, and neurons were incubated with the primary antibody for 1 hour at RT. Neurons were washed three times with PBS and stained with fluorophore conjugated secondary antibodies in blocking buffer for 1 hour at RT. The samples were post-fixed with 4% PFA for 20 min.

### STORM imaging

The STORM setup was based on a Nikon Eclipse-Ti inverted microscope or Olympus IX71. 405-nm (OBIS 405-50C; Coherent) and 647-nm (F-04306-113; MPB Communications) lasers were introduced into the sample through the back port of the microscope. A translation stage allowed the laser beams to be shifted towards the edge of the objective so that the emerging light reached the sample at incidence angles slightly smaller than the critical angle of the glass-water interface, thus illuminating only the fluorophores within a few micrometers of the coverslip surface. T660LPXR (Chroma) was used as the dichroic mirror and an ET705/72M band-pass filter (Chroma) was used as the emission filter. For one-color three-dimensional (3D) STORM imaging, a cylindrical lens was inserted between the microscope side port and the EMCCD camera (Andor iXon, DU-897E-CSO-#BV, Andor Technology) so that images of single molecules were elongated in *x* and *y* for molecules on the proximal and distal sides of the focal plane (relative to the objective), respectively, and the ellipticity of the single-molecule images were used to determine the z position of the molecules ^22^. For two-color two-dimensional STORM imaging, a multichannel imaging system (Photometrics, QV2) was inserted between microscope body and the EMCCD camera. Using the QV2 system, the emission light was split by T685LPXR (Chroma) into the two channels for the two-color STORM imaging acquisition. As a result, signals from CF680 and Alexa 647 were split into two color-channels, and the fluorophores were identified by measuring the intensity ratio of each single molecule in two channels ^64, 65^.

The sample was imaged in PBS buffer containing 100 mM cysteamine (Sigma), 5% glucose (Sigma), 0.8 mg/mL glucose oxidase (Sigma), and 40 µg/mL catalase (Roche Applied Science). During imaging, continuous illumination of 647-nm laser (∼2 kW/cm^2^) was used to excite fluorescence from Alexa Flour 647 or CF680 molecules. Continuous illumination of the 405-nm laser was used to reactivate the fluorophores to the emitting state. The intensity of the activation laser (0-1 W/cm^2^) was adjusted during image acquisition so that at any given instant, only a small fraction of the fluorophores in the sample were in the emitting state, such that their images are optically resolvable and their positions can be determined from their images.

A typical STORM image was generated from a sequence of about 25,000∼40,000 image frames at a frame rate of 60 Hz. The recorded single-color 3D STORM movie was analyzed according to previously described methods ^21, 22^. Briefly, the centroid positions and ellipticities of the single-molecule images provided lateral and axial positions of each activated fluorescent molecule, respectively ^22^. Super-resolution images were reconstructed from the molecular coordinates by depicting each location as a 2D Gaussian peak. For two-color STORM analysis, each channel was analyzed using the same method as the single-color STORM analysis. Two channels were then aligned, and the intensity ratio of each aligned fluorophore between two channels was compared and assigned to the correct color ^64, 65^.

### GO term enrichment analyses

The GO term enrichment analyses were performed using the Database for Annotation, Visualization, and Integrated Discovery (DAVID, https://david.ncifcrf.gov/) ^66^. The official gene symbols of the candidate MPS-interacting proteins were submitted to DAVID, and the enriched GO terms with *p*-value ≤ 0.05 were considered as significant. For GO term enrichment analysis of biological process (BP) category, GO BP terms at level 3 (GO_BP_3) were used, and the top nine enriched clusters (Fig. 1d) were determined by automated functional annotation clustering based on enriched GO_BP_3 terms. For GO term enrichment analysis of molecular function (MF) category, GO MF terms at level 4 (GO_MF_4) were determined for the candidate MPS-interacting proteins (Supplementary Fig. 1b), and GO MF terms at levels 4 and 5 (GO_MF_4 and GO_MF_5) were used for the candidate MPS-interacting transmembrane proteins (Fig. 4a).

### Autocorrelation analysis and cross-correlation analysis

For 1D autocorrelation analysis of each molecule, the signals in the STORM images were projected to the longitudinal axis of the axon segments, and autocorrelation functions were calculated from these 1D projected signals. For the average autocorrelation function, the autocorrelation curves from 50-100 randomly selected axon segments (∼ 2 µm in length) were averaged for each condition. The average 1D autocorrelation amplitude was defined as the difference between the first peak (at ∼190 nm) and the average of the two first valleys (at ∼95 nm and ∼285 nm) of the average 1D autocorrelation curve ^4, 10^. The average period (or spacing) of the MPS structure was defined as the first peak position of 1D average autocorrelation function. For 1D cross-correlation analysis between two molecules (NMIIA/B and βII spectrin), the signals in the two color channels of the STORM images were each projected to the longitudinal axis of the axon segments, and a cross-correlation function was calculated from these projected signals from the two color channels for axon segment, and then averaged over 50∼100 randomly selected axonal segments (∼ 2 µm in length) for each condition.

### Axon-axon and axon-dendrite bundling analyses

For axon-axon bundling, the morphology of immunostained axons in the field of view of conventional fluorescence images were determined using Otsu’s threshold, and the average width of axon-axon bundles were calculated using the ImageJ plugin DiameterJ, a previously validated nanofiber diameter characterization tool to find the central axis and average diameter of tubular structures in an image ^67^. For axon-dendrite interaction, the degree of axon-dendrite bundling was calculated as the fraction of the dendritic length that shows overlapping axon and dendrite signals. The synapse density was obtained by calculating the number of coupled pre-synaptic and post-synaptic markers divided by total dendrite area.

### Data availability

All data supporting the findings of this study are included in this published article and its Supplementary Information. Any other data and material will be made available upon reasonable request.

### Code availability

Custom Python and MATLab codes for image acquisition and STORM analysis are available at https://github.com/ZhuangLab. Custom MATLab codes for the two-color STORM data analysis, autocorrelation and cross-correlation analysis are available at: https://github.com/boranhan/MPS.

## Acknowledgements

We thank Dr. Jiang He for help with plasmid construction, Dr. Ke Xu for providing the software for ratiometric two-color STORM analysis, and Dr. Bogdan Budnik and Ms. Renee Robinson for help with mass-spectrometry analyses. This work is supported in part by the National Institutes of Health. R.Z. is an HHMI Fellow of the Life Sciences Research Foundation. X.Z. is a Howard Hughes Medical Institute investigator.

## Author contributions

R.Z., B.H. and X.Z. designed the experiments with input from A.H.C. and V.M.F.. R.Z., B.H., R.N, Y.L, E.H, and C.X. performed the experiments. B.H. and R.Z. performed data analysis. R.Z., B.H., and X.Z. interpreted the data and wrote the manuscript with input from A.H.C., V.M.F and the other authors.

## Competing interests

The authors declare no competing interests.

## Supplementary Information

### Supplementary figures

**Supplementary Figure 1:**
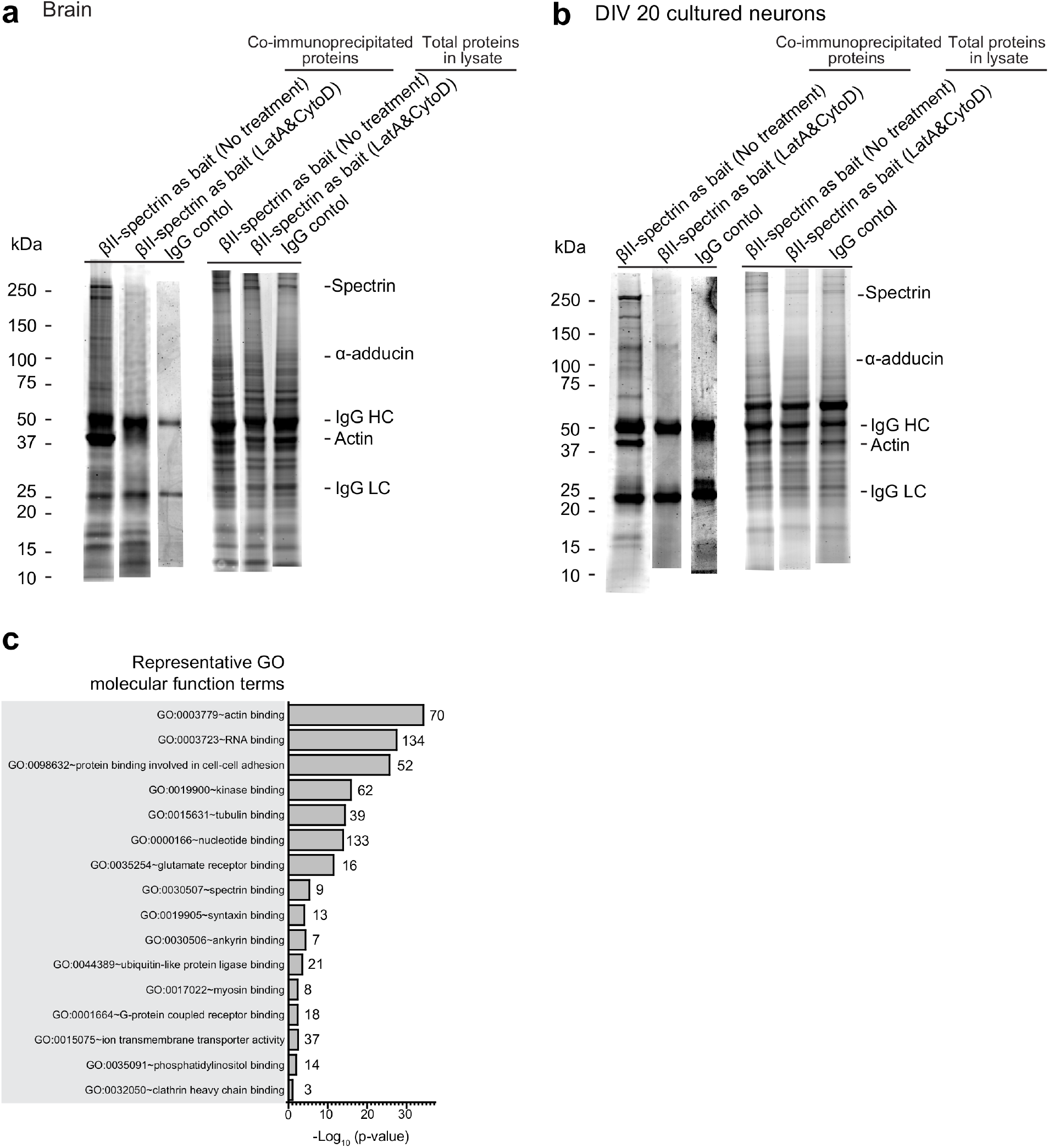
SDS-PAGE images of co-immunoprecipitated proteins from adult mouse brains and cultured mouse neurons and GO term analyses in the molecular function category of candidate MPS-interacting proteins. **a** SYPRO Red-stained SDS-PAGE images of co-immunoprecipitated proteins from the adult mouse brain using the βII-spectrin antibody (left most lane) and an irrelevant antibody as the control (third left lane). To perform the co-IP experiments under conditions where the MPS was disrupted, F-actin disrupting drugs (LatA and CytoD) were added to the lysate prior to incubating the lysate with the beads coated with the βII-spectrin antibody (second left lane). Also shown are the corresponding SYPRO Red-stained SDS-PAGE images of total cell proteins in the lysates from the adult mouse brain (right three lanes). **b** Same as (**a**) but for cultured mouse hippocampal neurons (DIV 20) instead of adult mouse brain. **c** Enriched GO molecular function (MF) terms of the 515 candidate MPS-interacting proteins. Enriched GO MF terms were generated using the DAVID 6.8 platform, and representative GO MF terms were shown with the corresponding *p*-values (bars) and the protein numbers (next to the bars) for each enriched GO MF term.

**Supplementary Figure 2:**
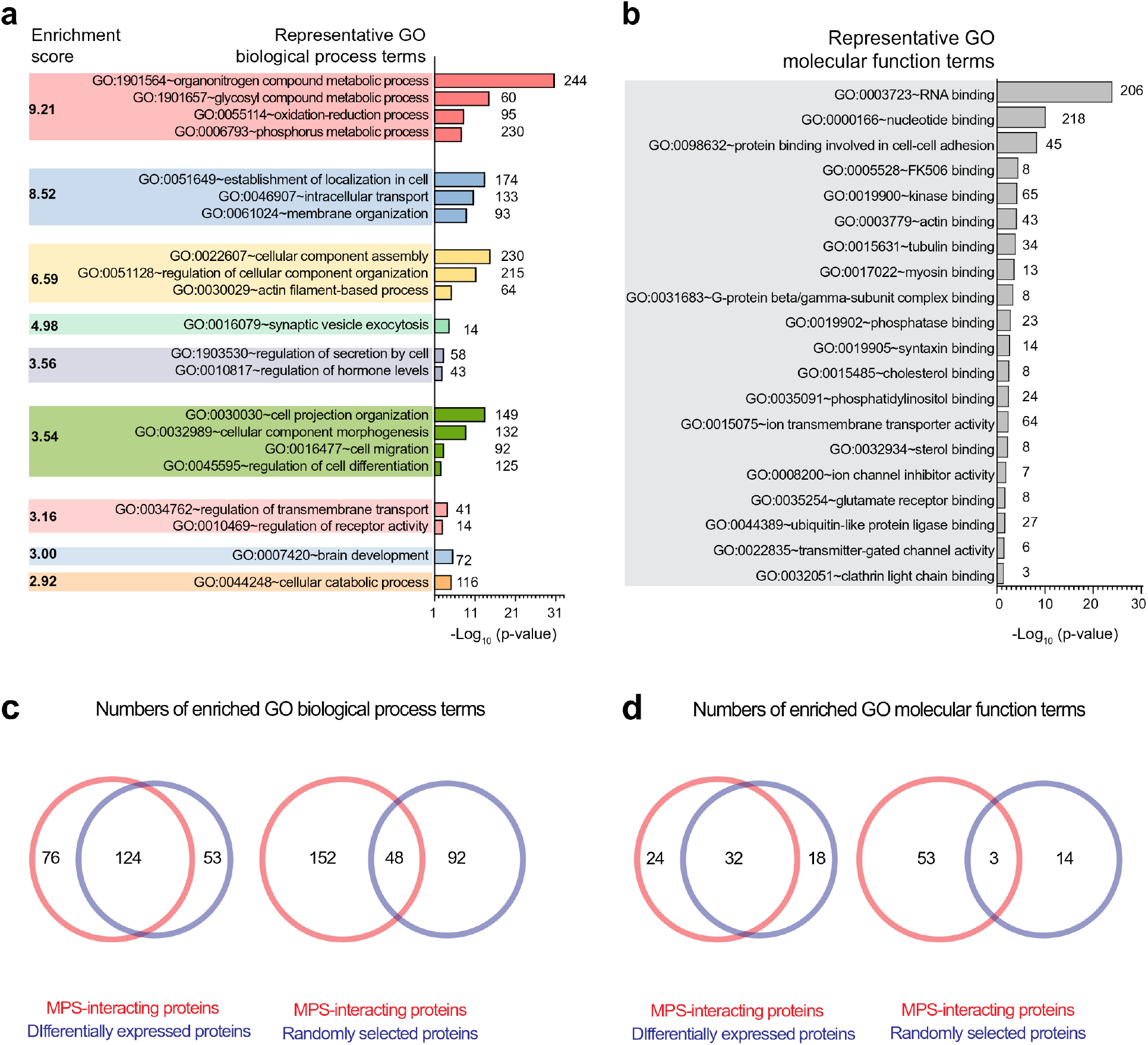
GO term analyses for the differentially expressed proteins in MPS-disrupted neurons. **a** Functional annotation clustering of the enriched GO terms in the biological process (BP) category for the 1204 differentially expressed proteins in βII-spectrin knockdown neurons. Enriched GO BP terms were generated and clustered using the DAVID 6.8 platform with the enrichment score listed for each color-coded GO term cluster. Representative GO BP terms for each GO term cluster were shown. The corresponding *p*-values (bars) and the protein numbers (next to the bars) for each enriched GO BP term are shown on the right. **b** Enriched GO molecular function (MF) terms of the 1204 differentially expressed proteins in βII-spectrin knockdown neurons. Enriched GO MF terms were generated using the DAVID 6.8 platform, and representative GO MF terms were shown with the corresponding *p*-values (bars) and the protein numbers (next to the bars) for each enriched GO MF term. **c** Venn diagrams showing the overlap between enriched GO BP terms of the 515 candidate MPS-interacting proteins and enriched GO BP terms of the 1204 differentially expressed proteins (left), in comparison with the overlap between enriched GO BP terms of the 515 candidate MPS-interacting proteins and enriched GO BP terms of 1204 randomly selected genes from the mouse genome (right). **d** Venn diagrams showing the overlap between enriched GO MF terms of the 515 candidate MPS-interacting proteins and enriched GO MF terms of the 1204 differentially expressed proteins (left), in comparison with the overlap between enriched GO MF terms of the 515 candidate MPS-interacting proteins and enriched GO MF terms of 1204 randomly selected genes from the mouse genome (right).

**Supplementary Figure 3:**
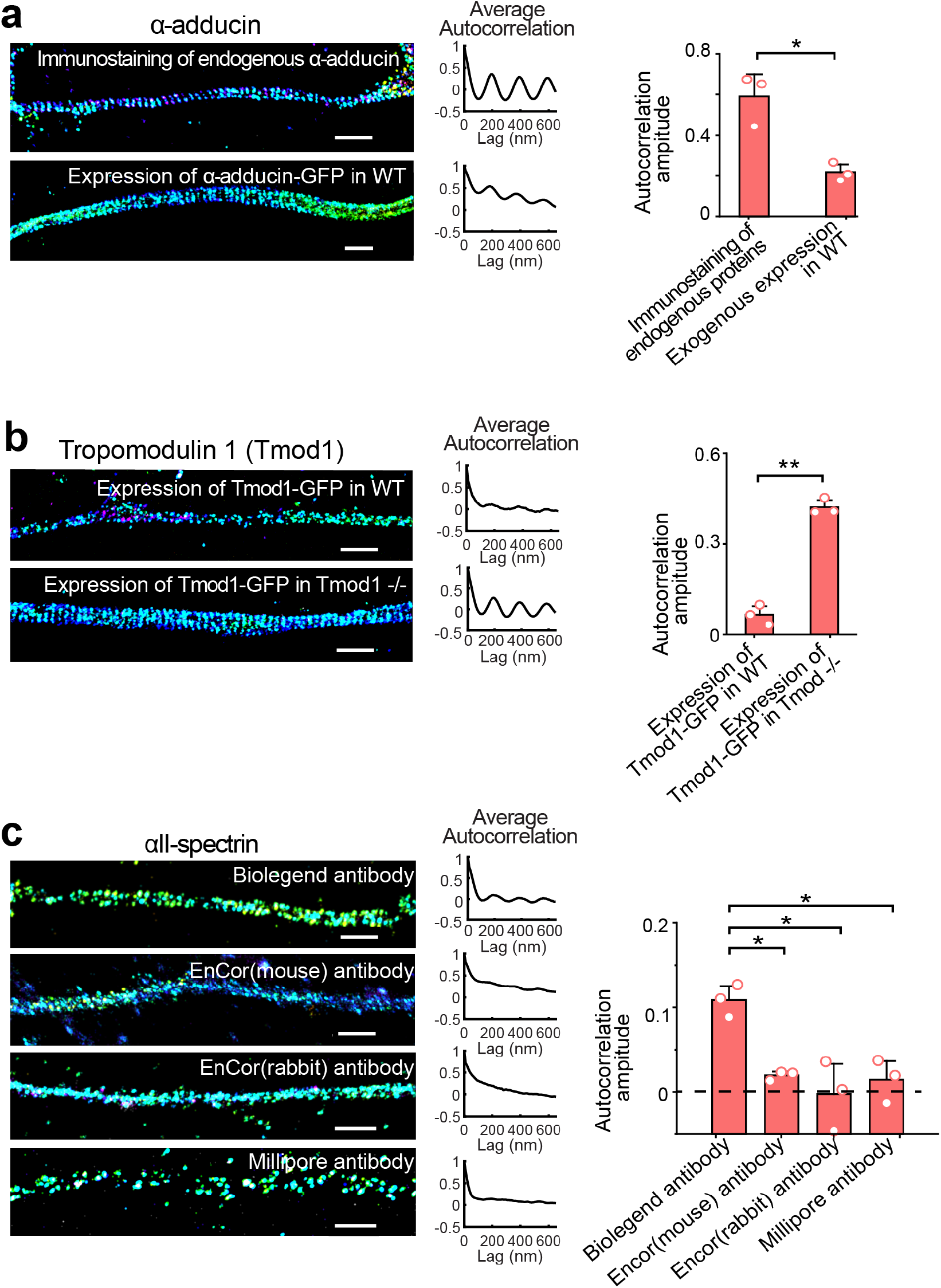
Super-resolution imaging of MPS-interaction proteins using different labeling strategies. **a** Left: 3D STORM images of α-adducin in axons of cultured hippocampal neurons labeled using two different labeling strategies. In the first strategy, α-adducin was immunostained with the antibody against α-adducin (top). In the second strategy, α-adducin was labeled using moderate expression of GFP-tagged α-adducin through low-titer lentiviral transfection, followed by immunolabeling using anti-GFP antibody. Middle: Average 1D autocorrelation functions of α-adducin over many randomly selected axon regions for the two labeling strategies. One-dimensional (1D) autocorrelation function of the imaged molecule in any given axon region was calculated from the signals of the molecule projected to the longitudinal axis of the axon segment. The 1D autocorrelation function was then averaged over many axon segments. Right: Average 1D auto-correlation amplitudes for the two labeling strategies. The average 1D auto-correlation amplitude was defined as the difference between the first peak (at ∼190 nm) and the average of the two first valleys (at ∼95 nm and ∼285 nm, respectively) of the average 1D auto-correlation function^4^. * indicates *p* < 0.05 (unpaired student’s t-test). Scale bars: 1 µm. **b** Same as (**a**) but for tropomodulin 1 in the axons of neurons labeled using the second strategy (GFP fusion protein), with lentiviral transfection performed on either WT neurons (top) or tropomodulin 1 knockout neurons (bottom). * indicates *p* < 0.05 (unpaired student’s t-test). Scale bars: 1 µm. **c** Same as (**a**) but for αII-spectrin in the axons of neurons labeled using the first strategy (immunolabeling of endogenous proteins), with four different antibodies against αII-spectrin. * indicates *p* < 0.05 (unpaired student’s t-test). Scale bars: 1 µm.

**Supplementary Figure 4:**
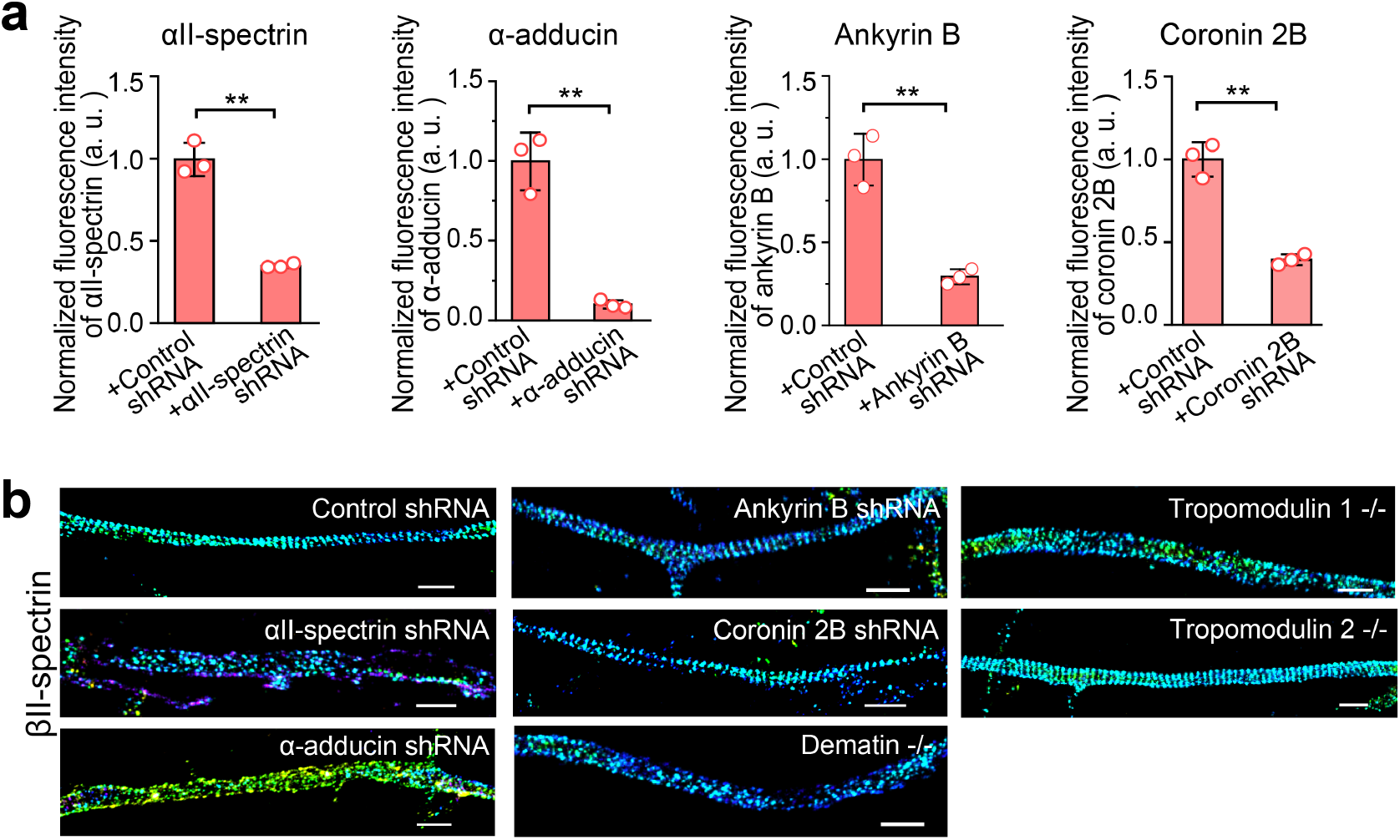
Determination of proteins essential for MPS formation and maintenance by gene knockdown and knockout. **a** Average fluorescence intensities of αII-spectrin, α-adducin, ankyrin B, and coronin 2B in the axons of neurons transfected with adenoviruses expressing scrambled (control) shRNA or the shRNA against the corresponding protein, showing the knockdown of αII-spectrin, α-adducin, ankyrin B, and coronin 2B. Data are mean ± s.e.m. m (*n* = 3 biological replicates; 10-20 imaged regions were examined per condition). ** indicates *p* < 0.005 (unpaired student’s t-test). **b** 3D STORM images of βII-spectrin in axon segments of neurons treated with control shRNA, αII-spectrin shRNA, α-adducin shRNA, ankyrin B shRNA, coronin 2B shRNA, and of neurons cultured from dematin, tropomodulin 1, and tropomodulin 2 knockout mice. Scale bars: 1 µm

**Supplementary Figure 5:**
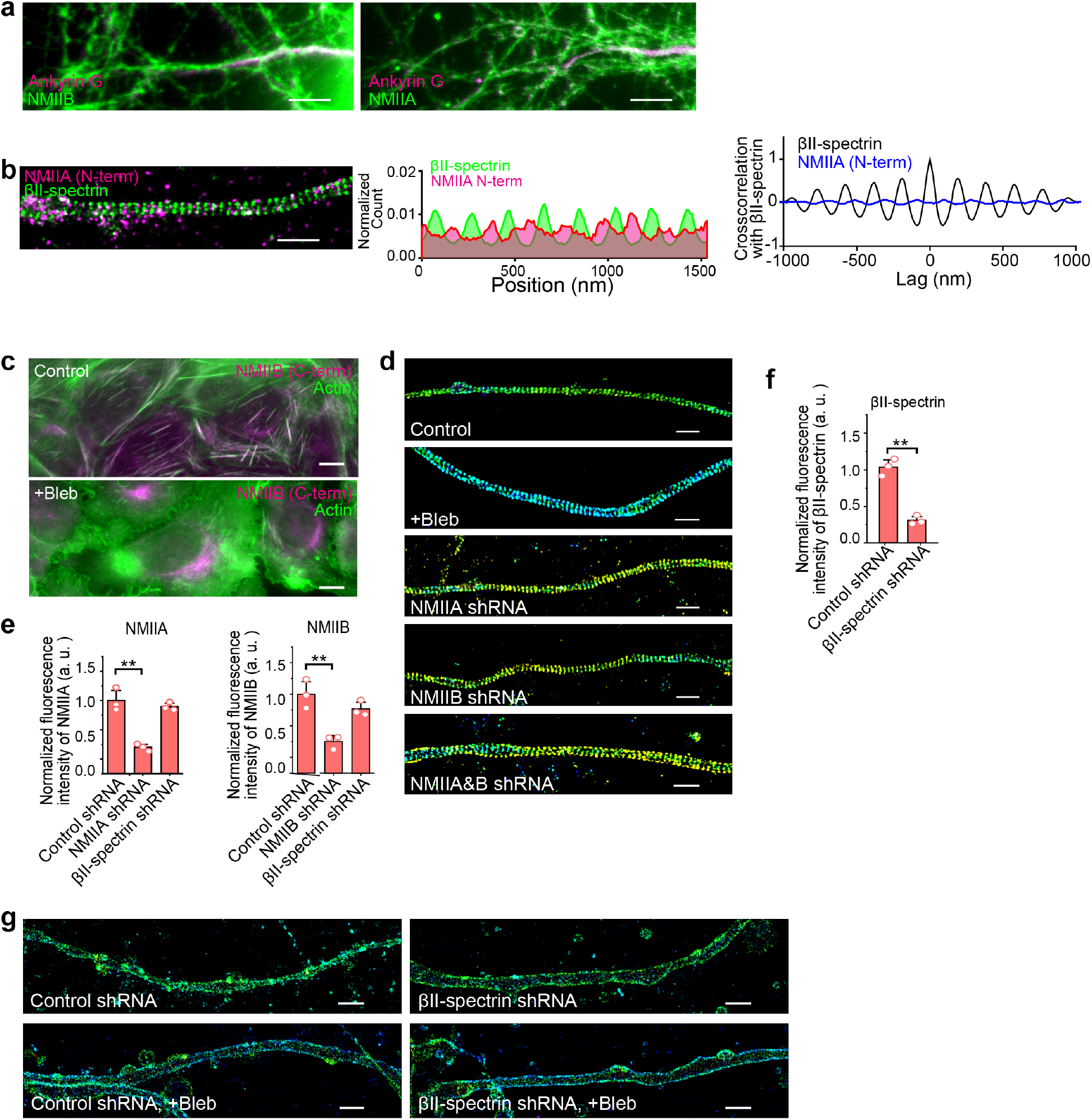
Spatial distribution of NMIIB and NMIIA in axons and effects of NMII activity inhibition on the MPS. **a** Left: Conventional fluorescence image of NMIIB (green) and the AIS marker ankyrin G (magenta) in axons of cultured neurons showing that NMIIB is expressed not only in the AIS but also in axon regions outside the AIS. Right: Similar to the left panel but for NMIIA instead of NMIIB. Scale bars: 10 µm. **b** Left: Two-color STORM image of βII-spectrin (C-terminus, green) and NMIIA (N-terminus, magenta) in the axons. Middle: Average 1D distribution of βII-spectrin (C-terminus, green) and NMIIA (N-terminus, magenta) signals projected to the longitudinal axon axis, derived from many axon regions. The axonal regions were aligned based on the phase of the βII-spectrin distribution for each analyzed region and the average 1D distributions were then calculated for both βII-spectrin and NMIIA. Right: Average 1D cross-correlation function between the distributions of βII-spectrin (C-terminus) and NMIIA (N-terminus) (blue). Average 1D auto-correlation function of βII-spectrin distribution (black) is reproduced from Fig. 3D as a reference. Scale bars: 1 µm. **c** Conventional fluorescence images of actin (green) and NMIIB (C-terminus, magenta) in untreated U2OS cells (top) and U2OS cells treated with blebbistatin (Bleb) for 2 hours (bottom). Actin was stained with phalloidin. Scale bars: 10 µm. **d** 3D STORM images of βII-spectrin in the axons of control untreated neurons, neurons treated with Bleb, and neurons treated with NMIIA heavy chain shRNA, NMIIB heavy chain shRNA or both shRNAs. Scale bars: 1 µm. **e** Average fluorescence intensities of NMIIA and NMIIB in the axons of neurons transfected with adenoviruses expressing shRNA against NMIIA heavy chain, NMIIB heavy chain, or βII-spectrin, in comparison with neurons transfected with adenoviruses expressing scrambled (control) shRNA. Data are mean ± s.e.m. (*n* = 3 biological replicates for each condition; for each replicate, 10-20 imaged regions were examined). ** indicates *p* < 0.005 (unpaired student’s t-test). **f** Average fluorescence intensities of βII-spectrin in the axons of neurons transfected with adenoviruses expressing scrambled (control) shRNA and βII-spectrin shRNA. Data are mean ± s.e.m. (*n* = 3 biological replicates for each condition; for each replicate, 10-20 imaged regions were examined). ** indicates *p* < 0.005 (unpaired student’s t-test). **g** 3D STORM images of the cholera toxin B (CTB)-stained axons of neurons treated with control (scramble) shRNA, neurons treated with control shRNA and Bleb, neurons treated with βII-spectrin shRNA, and neurons treated with βII-spectrin shRNA and Bleb. Scale bars: 1 µm.

**Supplementary Figure 6:**
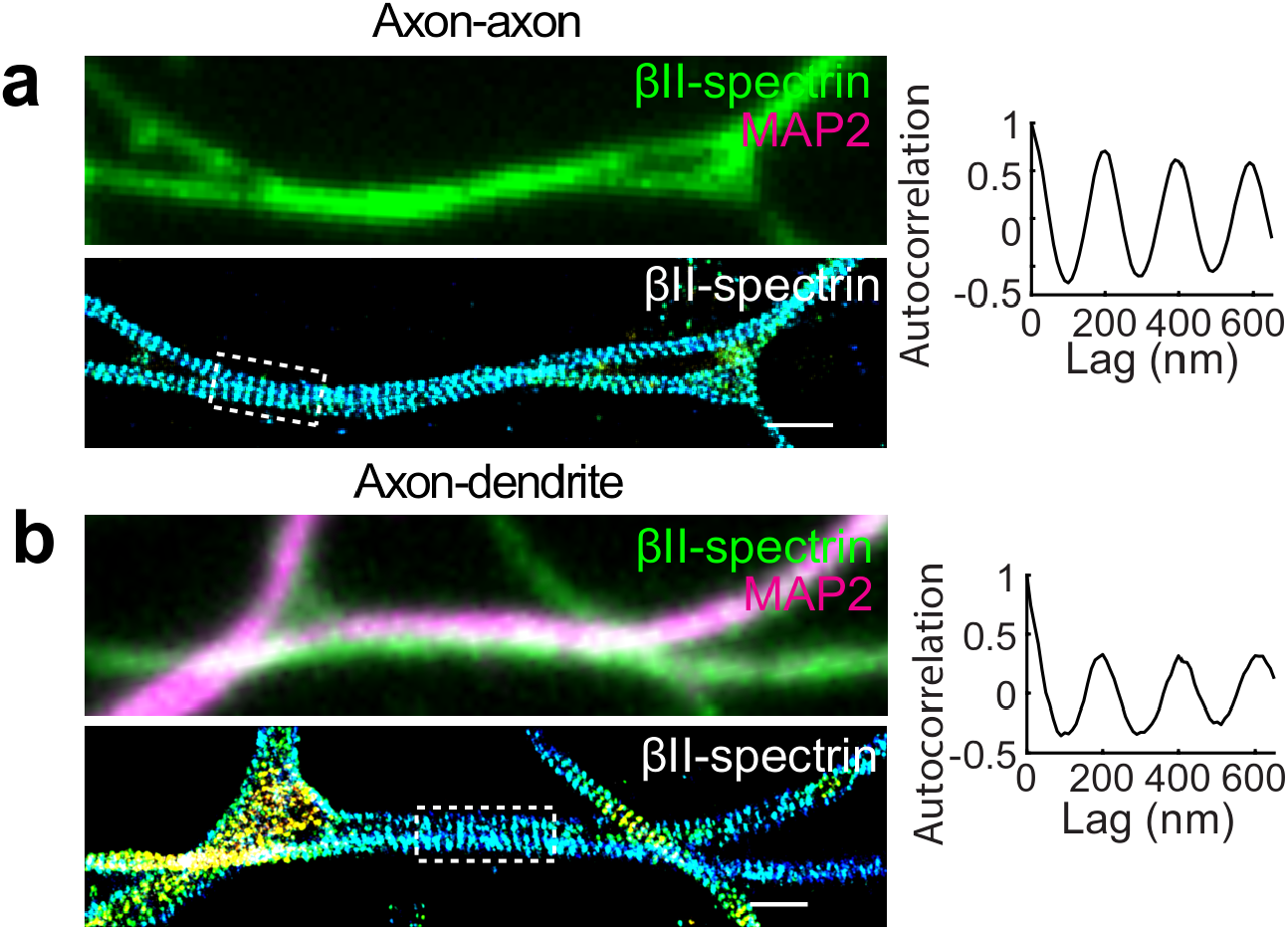
MPS alignment between abutting neurites. **a-b** Top left: conventional fluorescence images of cultured neurons immunostained for βII-spectrin (green) and MAP2 (magenta). Bottom left: 3D STORM images of βII-spectrin in the same region as shown in the top left panel, exhibiting MPS alignment at the axon-axon (**a**) and axon-dendrite (**b**) contact sites, exemplified by regions in the dashed boxes. Right: 1D autocorrelation function of the boxed regions in the bottom left panels. Scale bars: 1 µm.

**Supplementary Figure 7:**
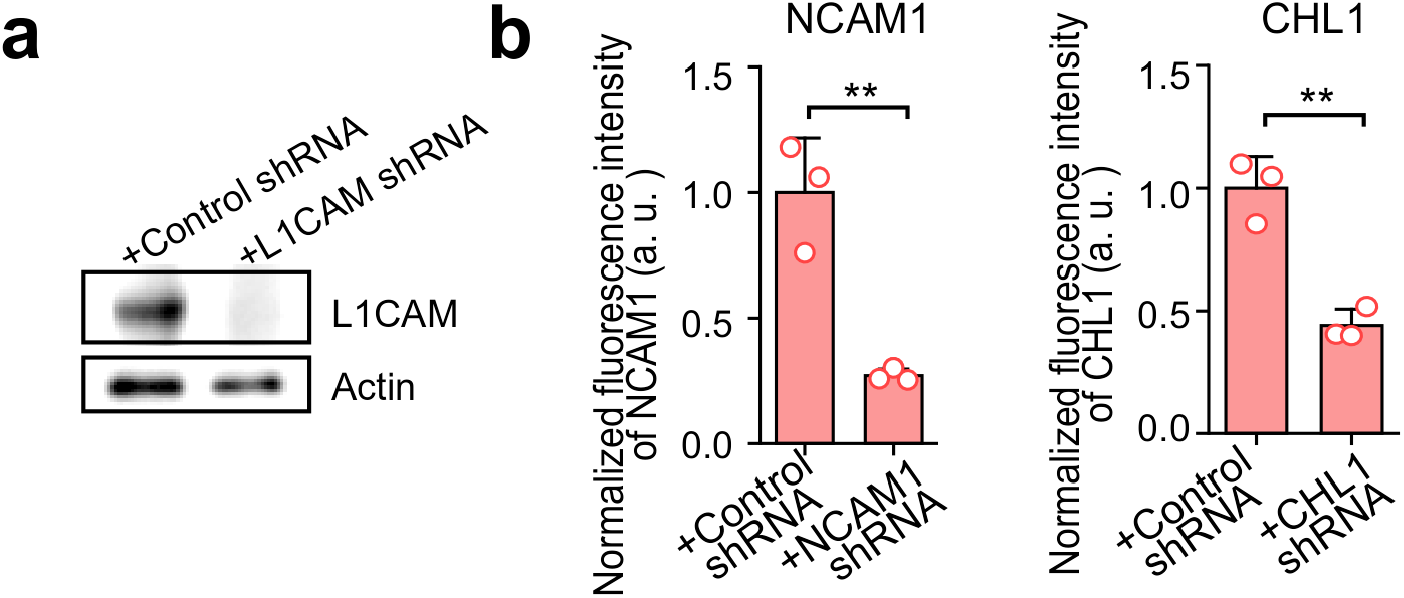
Knockdown of cell adhesion molecules. **a** Western blots showing the knockdown of L1CAM in the cultured neurons transfected with adenoviruses expressing L1CAM shRNA in comparison with neurons transfected with adenoviruses expressing control shRNA. **b** Average fluorescence intensities of NCAM1 and CHL1 in the axons of neurons transfected with adenoviruses expressing scrambled (control) shRNA or the shRNA against the corresponding protein (NCAM1 and CHL1). Data are mean ± s.e.m. (*n* = 3 biological replicates; 10-20 imaged regions were examined per condition). ** indicates *p* < 0.005 (unpaired student’s t-test).

**Supplementary Figure 8:**
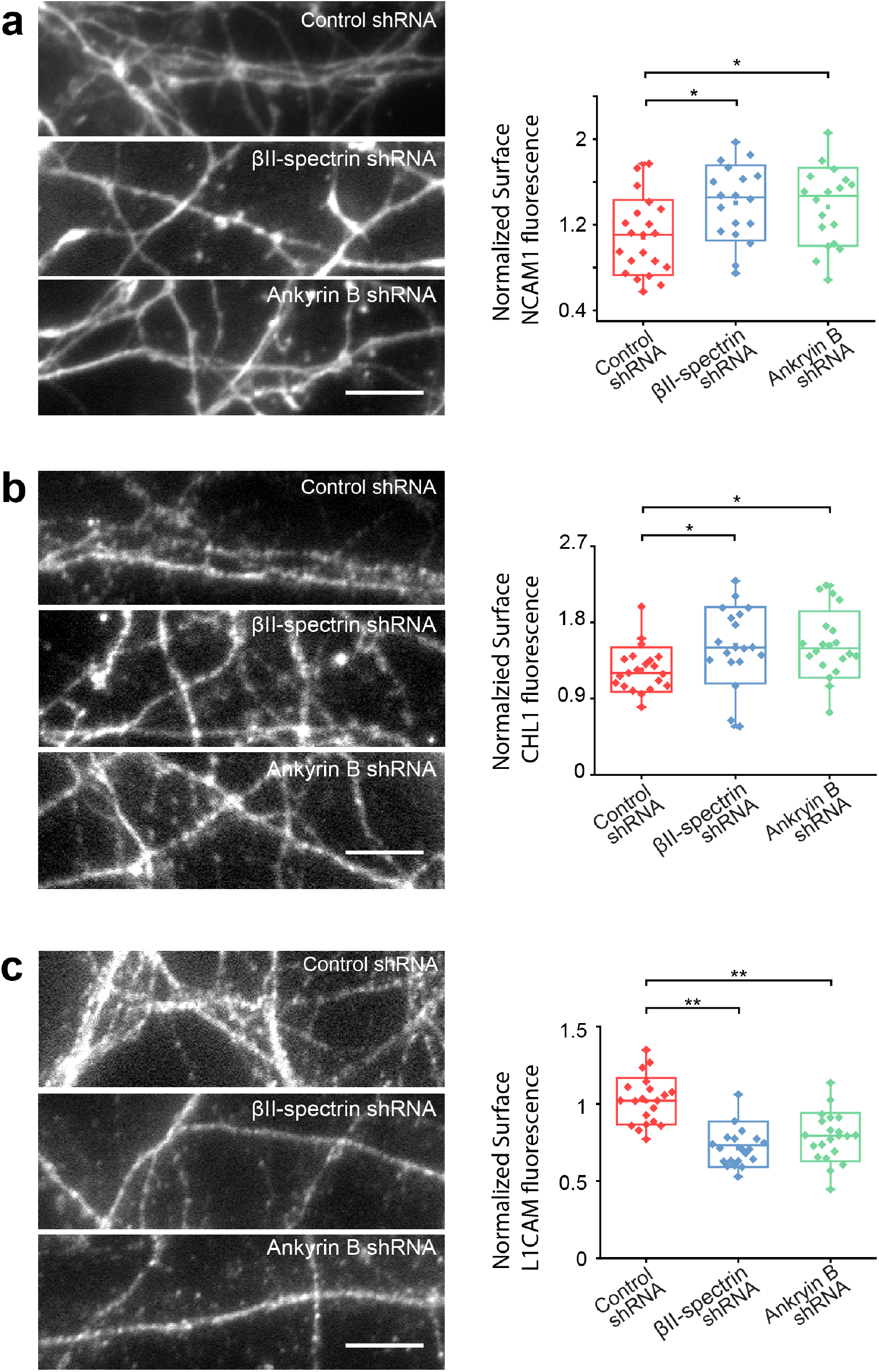
Cell-surface expression levels of NCAM1, CHL1 and L1CAM in control, βII-spectrin knockdown and ankyrin B knockdown neurons. **a** Left: Fluorescence images of cell-surface NCAM1 for the neurons transfected with adenoviruses expressing scrambled (control) shRNA (top panel), βII-spectrin shRNA (middle panel) and ankyrin B shRNA (bottom panel). Right: Box plots of the cell-surface NCAM1 expression levels at the neurites for neurons transfected with adenoviruses expressing control shRNA (red), βII-spectrin shRNA (blue) and ankyrin B shRNA (green). * indicates *p* < 0.05 and ** indicates *p* < 0.005 (unpaired student’s t-test). The line in the middle of the box indicates the median value; the lower and upper bounds of the box indicate 25^th^ and 75^th^ percentile; the whiskers indicate the minimum and maximum. **b** Same as (**a**) but for the cell-surface expression levels of CHL1 instead of NCAM1. **c** Same as (**a**) but for the cell-surface expression levels of L1CAM instead of NCAM1.

## Notes

### Competing Interest Statement

The authors have declared no competing interest.

